# Structures of muscle-type nicotinic acetylcholine receptor with α-conotoxins reveal determinants of receptor-subtype specificity

**DOI:** 10.64898/2026.01.15.699711

**Authors:** Michael J. Capper, Oscar A. Shepperson, Charlie Holdship, Oliver J. Melling, Nicola Wade, Michael A. Malone, Kirsty I.M. Arnott, Leonie Windeln, Steven Turner, Charlotte Whitmore, Tiffany D. Morcom, Josephine A. Connah, Christopher M. Timperley, Jeremy G. Frey, A. Christopher Green, Jonathan W. Essex, Jesko Köhnke, Andrew G. Jamieson

## Abstract

α-Conotoxins (α-CTX) are disulfide-rich peptide antagonists with exceptional potency and subtype selectivity across nicotinic acetylcholine receptors (nAChRs), making them promising leads for therapeutic development. The molecular basis of their specificity has remained unresolved for decades, limiting rational drug design. Here, we present the first high-resolution cryo-EM structures of full-length muscle-type nAChR bound to three distinct α-CTXs, revealing their precise binding modes at both acetylcholine binding sites. Our integrated approach, combining structural biology, pharmacological profiling, and computational hydration mapping, uncovers conserved pharmacophore features, water-mediated interactions, and subunit-specific determinants that govern potency and selectivity. These findings provide a long-sought molecular framework for α-conotoxin recognition and inhibition, addressing a critical gap required to realise the full potential of conotoxins as lead compounds in drug discovery.

## INTRODUCTION

Cone snails (*Conus sp.*) produce venom composed of diverse, pharmacologically active peptides to hunt and paralyse prey for feeding^1^. Central to this venom are short, disulfide-rich conotoxin peptides, that target a broad spectrum of receptors and channels, including ligand-gated ion channels, voltage-gated ion channels, G protein-coupled receptors (GPCRs) and membrane transporters^2^. The precise molecular determinants of specificity and potency have remained elusive, which has obstructed an understanding of the conotoxin mechanism of action and their rational development as pharmacological tools and therapeutics.

Each species of cone snail is estimated by transcriptomic and proteomic analyses to produce as many as 200 unique conotoxins. Collectively this contributes to a global total of approximately 80,000 distinct molecules^3,4^, of which only 10% have been experimentally characterised and catalogued in the ConoServer database^5^. This represents an untapped wealth of potentially therapeutic cysteine-rich cyclic peptide scaffolds, as exemplified by the synthetic conotoxin Ziconotide^TM^, an approved treatment for chronic pain^6^.

Each polycyclic conotoxin fold is constrained by disulfide bonds that are essential for both function and stability^7^. The pattern of cysteines in the peptide sequence (e.g. C_I_C_II_-C_III_-C_IV_; as defined by locations of cysteines and loops), and the folded disulfide connectivity, form conotoxin cysteine frameworks into which the peptides can be grouped. The frameworks can be further subdivided by the number of residues between cysteines (3/5, 4/7 etc.). In addition, each toxin is classified according to its physiological target and activity, annotated by a letter of the Greek alphabet, i.e. “α” conotoxins are antagonists of nicotinic acetylcholine receptors (nAChR), whilst “ω” conotoxins are antagonists of voltage-gated calcium channels^2^. The first conotoxin to be purified from cone snail venom was α-GI, which is particularly potent against human muscle-type receptors^8^. Envenomation results in nAChR antagonism and prevents the depolarisation of the post-synaptic membrane and subsequent muscle contraction^9,10^.

nAChRs are a diverse family of pentameric ligand-gated ion channels with extensive roles in addiction, depression, chronic pain and neurological diseases^11,12^. There are 16 human subunit isoforms that combine to yield functional homo- or hetero-pentamers with unique pharmacology. These functional pentamers contain at least one α-subunit, which is the principal face of the acetylcholine (ACh) binding-site formed with the complementary face of a neighbouring subunit in the extracellular domain (ECD). Each functional pentameric receptor can contain multiple ACh binding sites, depending on the number of α-subunits. In humans, muscle-type nAChRs are comprised of subunits (α1)_2_β1δγ (foetal form), or subunits (α1)_2_β1δε (adult form). As a result, they contain two non-equivalent ACh binding sites formed by different pairs of complementary subunits, the alpha-delta site (α-δ) and the alpha-gamma (α-γ) or alpha-epsilon site (α-ε), corresponding to the foetal and adult receptor isoforms, respectively.

Previous structural studies of α-CTX binding have relied on the soluble nAChR surrogate acetylcholine binding protein (AChBP), a homolog that broadly mimics the fold of the nAChR extracellular domain^13^. While these data have provided valuable insights, its low sequence identity to the α1 subunit limits how well it captures the true architecture of heteromeric nAChRs. As a result, our molecular understanding of α-conotoxin inhibition at native receptors has remained incomplete. Unlike the well-studied three-finger toxins (3FTXs) such as α-bungarotoxin, which primarily bind to the principal α-subunit face, α-CTXs span both the principal and complementary subunits^14^. This distinct binding mode is thought to underpin their remarkable subtype selectivity.

In this study we employed cryo-EM to determine the structures of full-length, muscle-type *Torpedo* nAChR bound to three different chemically synthesised conotoxins of varying primary sequences, cysteine frameworks and pharmacological profiles. By integrating structural data with pharmacology, and computational modelling, key molecular drivers of nAChR inhibition by α-CTXs have been identified. Understanding the molecular basis of conotoxin binding to the nAChR can inform the design of novel conotoxin-based drugs with improved specificity and efficacy. This knowledge may also enable the development of potent conotoxin-blocking agents, facilitating new strategies to neutralise toxin activity, which could advance treatments for envenomation.

## RESULTS

### Structures of muscle-type nAChR bound to conotoxins

To explore α-CTX potency and selectivity for nAChR we synthesised three different α-CTX: α-GI, α-MI and α-SII (Figure 1A). Each of these α-CTXs are muscle-type selective but possess varying potencies. α-GI and α-MI adopt framework I, characterised by two disulfide bonds, whereas α-SII is unusual in incorporating three disulfide bonds, placing it in framework II^5^. The linear precursor peptides were synthesised by 9-fluorenylmethyloxycarbonyl solid-phase peptide synthesis (Fmoc-SPPS). Following cleavage from the peptidyl resin, the linear peptides underwent oxidation under thermodynamic conditions to yield the desired canonical isomers (Supplementary additional peptide methods). To determine α-CTX potency, we employed an ACh calcium response assay as previously demonstrated (Figure 1B)^15^. Potencies correlate with literature precedent at other muscle-type nAChR verifying purification of the correct isomers^16,17^.

**Figure 1.**
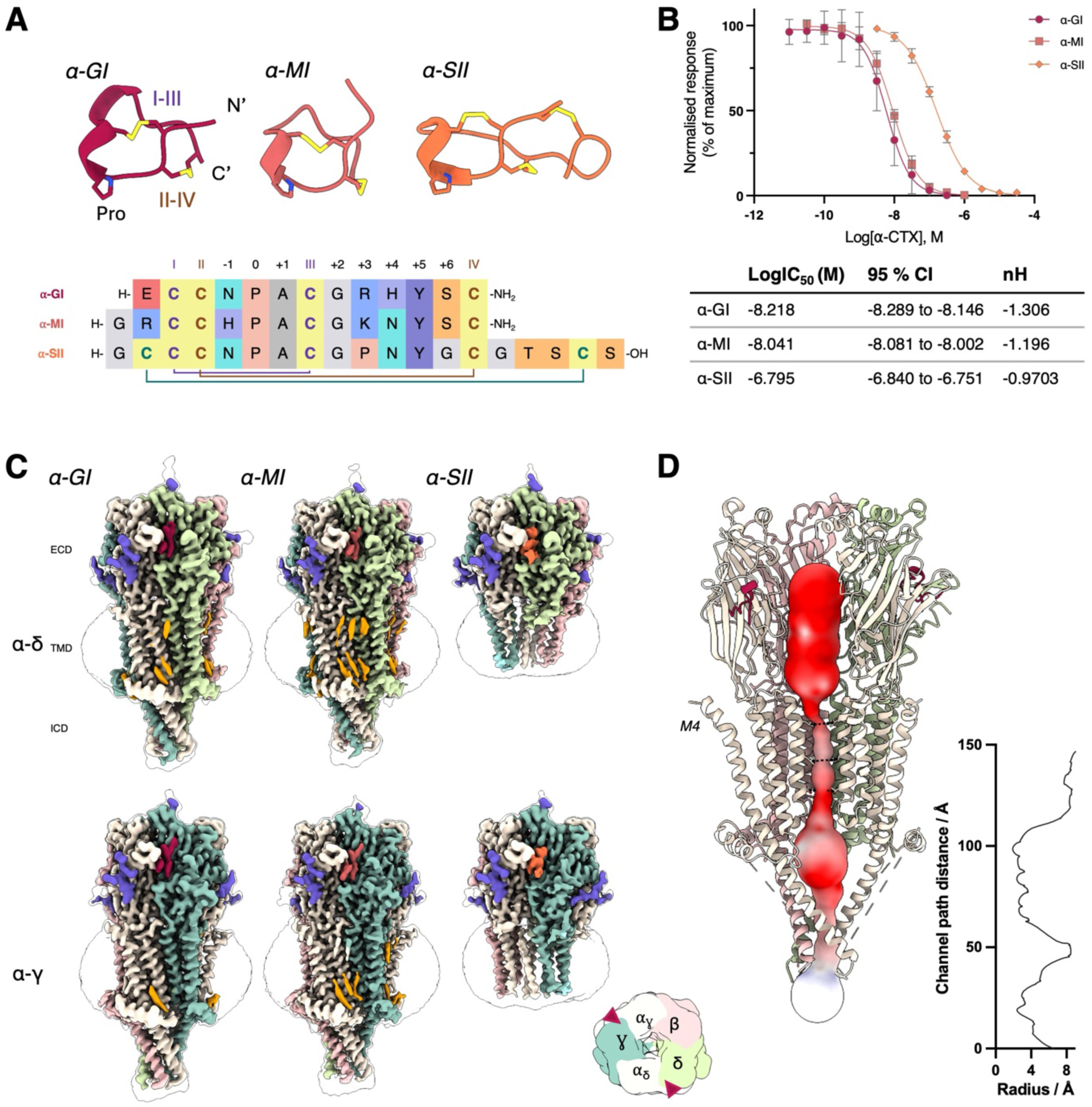
Conotoxin structures, activity and binding to muscle-type nAChR. (A) Aligned structures and sequences of the α-CTXs used in this study displaying the conserved proline residues and frameworks. Sequences display cysteine connectivity. (B) IC_50_ potencies and hill slopes (nH) for the toxins used in this study as determined by ACh calcium response data using α-CTX treated CN21ɣKO cells expressing human adult muscle type nAChR. All experiments were repeated in quadruplicate; error bars are standard deviations across all readings. (C) Cryo-EM volumes of detergent solubilised Torpedo muscle-type nAChR bound to each α-CTX (using the same colours as in Fig. 1A) at both the α-δ and α-γ binding sites). The receptor viewed from the top is shown as a schematic inset with toxin binding sites indicated by red triangles and subunit colours identified. Density for lipids and cholesterol are coloured orange and glycosylations in purple. (D) Path along the channel and diameter measurements taken through the representative α-GI bound nAChR structure identifies that the receptor is in the closed resting state.

Pacific electric ray (Torpedo*; Tetronarce californica*) muscle-type nAChR is comprised of subunits (α1)_2_β1δγ and has long been used as a model to understand nAChR structure and function. It corresponds to the foetal form of the human receptor and shares 75.3% sequence similarity and 60.4% sequence identity across all five subunits (Figure S1). We purified the receptor from *T. californica* tissue using a slight modification of established protocols (STAR methods)^18,19^, which resulted in high-resolution cryo-EM maps for subsequent model building (Figures 1C, S2-6, Table S1). We were able to attain high-resolution structures using detergent solubilised protein and thus avoided the need for nanodisc assembly. To determine the α-CTX-bound nAChR structures, we incubated purified receptor with the synthesised conotoxins prior to grid freezing. The overall structure of our α-CTX-bound *T. californica* nAChR closely aligns with earlier nAChR models^18,20^. When the complex is viewed from above, the five subunits α1-γ-α1-δ-β1 arrange clockwise to form the central pore (Figure 1C inset). The ECD was determined to particularly high-resolution with the α-CTX binding sites reaching local resolutions better than 2.5 Å supporting confident placement of key side chains and ordered water molecules (Figure S7). Bound cholesterol molecules and lipid molecules were identified in the transmembrane domain (TMD). The α1-subunit post-M4 helix, which moves considerably upon agonist binding, is present in the attached conformation alongside the ECD, which presents the receptor in its closed resting state as demonstrated by channel diameter analysis (Figure 1D)^21^. In the α-SII-bound receptor structure, high-resolution features were confided to the ECD, consistent with the absence of cholesterol during solubilisation and purification and highlighting the contribution of cholesterol to receptor stability.

### α-CTX binding mode

Each structure allowed for the unambiguous placement of an α-CTX at each of the two ACh binding sites (Figure 1C, Figure S7). Unlike the 3FTXs, such as bungarotoxin and the consensus short-chain α-neurotoxin (scNTX), the α-CTX binding sites overlap with that of ACh and are located beneath the C-loop of the principal face formed by the α1-subunit (Figure S8). Conotoxin binding therefore precludes ACh agonism of the muscle-type nAChR at the neuromuscular junction, preventing channel opening and depolarisation of the post synaptic membrane. This stops muscle contraction and results in flaccid paralysis^18,21,22^. Whilst 3FTX interactions are largely confined to the principal face of nAChR^20^, α-CTXs form extensive interactions with both the principal and complementary faces which we propose affords their selectivity for specific nAChR subtypes (Table S2).

To facilitate discussion of amino acid sequence features and the determinants of α-CTX activity, we have divided the conotoxin sequence at the central proline residue (defined as residue Pro-0). Non-cysteine residues *N-*terminal of Pro-0 have been assigned negative integers, while non-cysteine residues C-terminal of Pro-0 to the *C-*terminal have been assigned positive integers (Figure 1A).The α-CTXs adopt the same binding mode and orientation at both the α-δ and α-γ interfaces. In both cases, the N- and C-termini of the α-CTXs remain solvent exposed (Figures 2A and 2B). In addition to its role as a helix breaker at the centre of the α-CTX, Pro-0 forms extensive hydrophobic interactions within the aromatic cage of the choline binding site, formed by residues of the principal and complementary face (α1Y93/α1W149 and δW57/δL121 or γW55/γL119). Pro-0 also divides the α-CTX sequence into residues interacting with the principal face (*N-*terminal of Pro-0), and those interacting with the complementary face (*C-*terminal of Pro-0). The solvent facing region of each α-CTX is highly polar with a positive charge on the *N*-terminus and a C-terminal amide forming interactions with the highly charged acidic region of Loop F. This is consistent with the importance of *C-*terminal amidation for α-CTX activity^23^. ɑ-SII does not possess an amidated C-terminus but rather an additional disulfide constrained loop which points into bulk solvent and appears highly flexible in the cryo-EM structures which prevented accurate model building (Figure S9).

**Figure 2.**
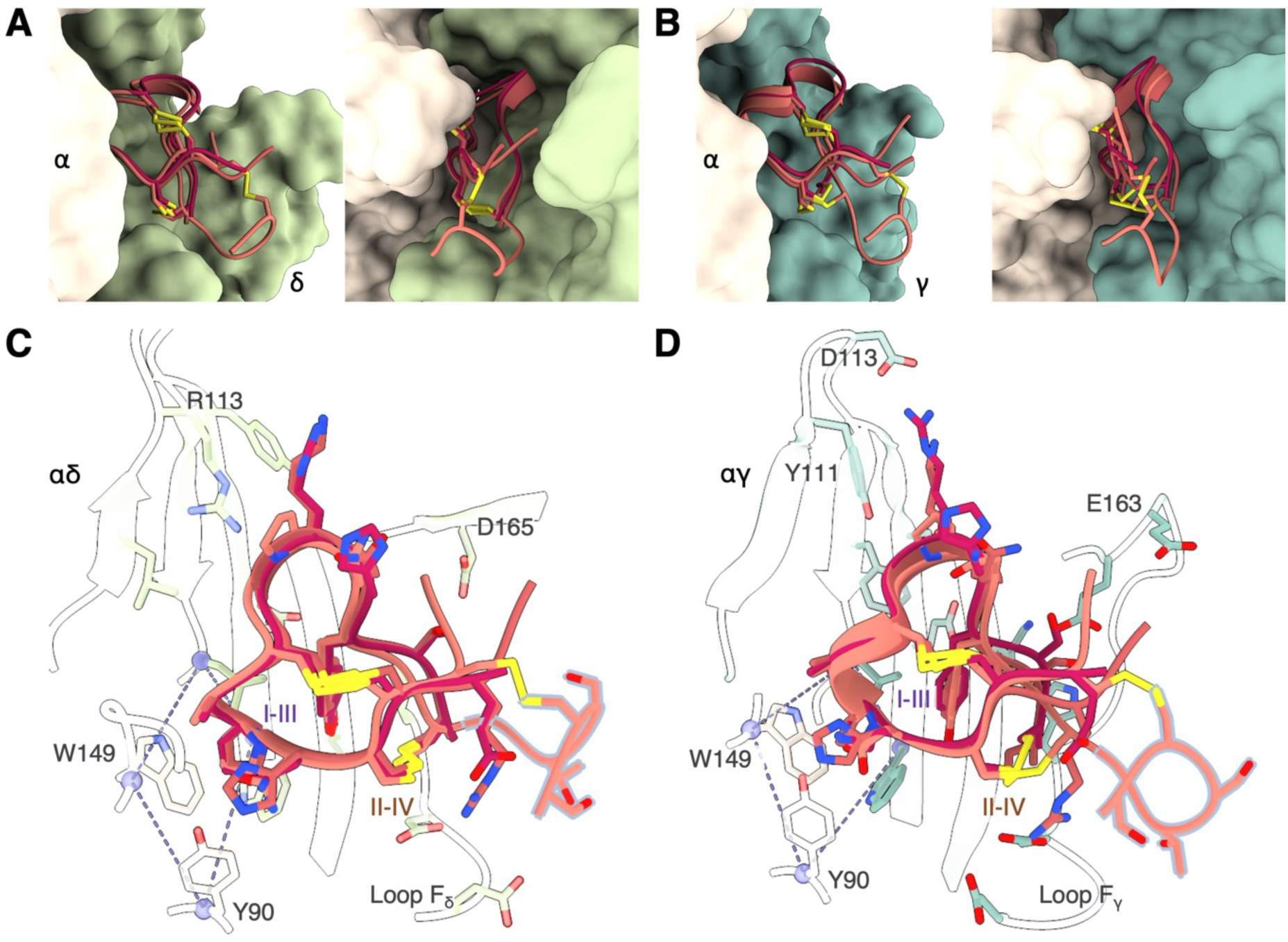
Conotoxin binding interactions at muscle-type nAChR. (A and B) Structures of the bound α-CTXs at the α-δ and α-γ sites respectively. Each image is rotated by 90° and loop C has been removed to facilitate viewing. (C and D) Overlay of each α-CTXs reveals overlapped bind modes possessing the same fold at the α-δ and α-γ sites respectively. Pro-0, Ala+1 overlap in each structure enclosed within the ACh binding aromatic cage (indicated by dashed lines). The *C-*terminal α-SII loop has been modelled into the unsharpened cryo-EM map (highlighted in grey).

The α-CTX disulfide bonds are resolved, although density is less well defined for the CysII-IV disulfide that projects toward the bulk solvent. The small loop disulfide bond (CysI-CysIII) of the conotoxin is oriented close to the vicinal disulfide bond of the principal face C-loop (αC192, αC193). The C-loop wraps around the α-CTX, facilitating tight binding and placing the α-CTX I-III disulfide bond next to the nAChR C-loop vicinal disulfide bond surrounded by hydrophobic residues α1Y190/α1Y198. In the Coulomb potential map, there is unambiguous density between nAChR C-loop αC192 and CysI of each conotoxin and the two cysteines can be fitted into the cryo-EM map such that an intermolecular disulfide bond is formed, and the I-III CTX disulfide is therefore broken. (Figure S10).

### α-δ α-CTX binding site

The torpedo muscle-type nicotinic acetylcholine receptor (nAChR) has a subunit composition of (α1)_2_β1δγ. This arrangement creates two distinct binding sites for α-CTX: one at the α-δ interface and another at the α-γ interface (Figures 2, S11 and S12, Tables S3 and S4). α-CTXs recognise the two agonist-binding interfaces with differing selectivity: α-GI and α-MI favour the *T. californica* α-γ site over the α-δ site, while α-SII binds both sites with comparable affinity^16,24–26^. The side-chain of the conserved α-CTX Tyr in the +5 position forms a tight interaction, locked between the conotoxin and hydrophobic residues δW57/δL121, whilst additionally forming a hydrogen bond via its hydroxyl group with the side-chain of δT38. The Ser at position +6 forms a hydrogen bond with the side-chain of δD165. The *N*-terminal Arg-2 of α-MI extends from the principal α face to the complementary δ face but does not appear to form a salt-bridge with δD180. Both α-GI and α-MI possess a cationic residue at the +3 position, which had previously been identified as a key driver of potency^27^. In our structures, the side-chain of this residue (Arg in α-GI and Lys in α-MI) extends across the ECD and appears to engage with δR113. Additionally, δR113 interacts with the α-CTX backbone through a water mediated hydrogen bond. The equivalent site in α-SII is shared by a Pro residue which precludes backbone interactions and correlates with reduced IC_50_ in the ACh calcium mobilisation assay (Figure 1B).

We have identified multiple conserved, ordered water molecules that strongly interact with the α-CTXs and facilitate binding (Figure S13). The carbonyl oxygens of residues +3 and +5 are hydrogen bonded to a network of water molecules that in turn form hydrogen bonds with receptor residues γK34, γE57 and γN59. The hydrogen bond donor at the -1 position in the α-CTX sequence forms a water mediated hydrogen bond through a trapped water to αY93 and the carbonyl of αL199. Three additional water molecules are positioned above the aromatic cage and interact with the carbonyl of Pro-0. To verify the inclusion of these waters in our model, molecular dynamics (MD) simulations were performed using a Monte Carlo enhanced sampling technique to facilitate fast convergence of the locations of occluded hydration sites. The locations of the water molecules around the conotoxins were clustered across the simulation frames.

Many of the high occupancy clusters were consistent across each of the α-CTXs in this study and overlay with water molecules built into the cryo-EM maps (Figure S14). The α face shows many hydration sites that are conserved across all three of the α-CTXs in both the cryo-EM density and water simulations. Three hydration sites located closer to the α-CTXs are only present with high occupancy when α-GI or α-SII are bound, and not when α-MI is bound. This discrepancy can be attributed to the presence of a His residue at the -1 position in α-MI, compared to an Asn in α-GI and α-SII. This His residue occupies the space of two of these hydration sites and results in a loss of potential hydrogen bonds that the water molecules can form. The simulations identify four hydration sites behind the conotoxins in the binding site that mediate interactions between both neighbouring subunits and the conotoxin residues. All four of these sites are seen with high occupancy and are conserved across all α-CTXs in this study. At the δ face the results are less clear, with the location and occupancies of the simulated water clusters varying across each structure. We attribute this to the fact that this interface is more solvent-exposed than at the α face meaning the hydration is more bulk-like and homogeneous.

### Binding differences at the α-γ site

The α-γ binding-site, though similar, presents a distinct environment due to differences between the δ and γ subunits. The α-CTX cationic residue Arg/Lys+3 extends to γY111 to form a cation-π interaction. In the case of α-GI, but not α-MI, Arg+4 forms an additional salt-bridge with γD113. Tyr+5 is again bound between the conotoxin Pro-0 residue and the hydrophobic residues γW55/γL119 and forms a weak hydrogen bond with the side-chain of γT36. α-CTX Ser+6 forms a hydrogen bond with the side-chain of γE163. E163 has a longer side-chain than the Asp residue at the α-δ interface, which results in a shorter hydrogen bond distance that likely strengthens the interaction. The *N-*terminal Arg-2, present in α-MI but not α-GI, orients across the solvent exposed face of the conotoxin to form a salt bridge with the side-chain of γD174, a residue that is not found at the α-δ site.

Water was added to the model as in the α-δ binding mode, and equivalent buried water molecules were observed to also interact with the carbonyl of Pro-0 (Figure S14). Comparable waters were also identified acting as hydrogen bond intermediates between the α-CTXs and the complementary face. Specifically, water molecules bound to the carbonyls at positions +3 and +5 also interact with γE57 and γK34 on the complementary face.

Analysis of the water clustering results from the MD simulations showed a similar picture to the α-δ binding site, with the His-1 residue in α-MI displacing some of the bound waters at the α face that are otherwise observed with the α-GI and α-SII conotoxins. Buried behind the α-CTXs, two conserved water molecules are observed instead of four. However, as with the α-δ site, both these hydration sites remain conserved and highly occupied. At the γ interface, the results are again less clear. Two hydration sites are observed that are conserved across all three structures, albeit with lower occupancies, and a number of other hydration sites are identified that are highly dependent on the specific α-CTX that is bound which may play a limited role in binding site selectivity.

### α-CTX affinity for human muscle-type nAChR

Adult human nAChR has high sequence similarity to that of *T. californica* (Figures 3A,B and S1). The corresponding human residues of the α-subunit that interact with the conotoxins in the *T. californica* are identical and the complementary subunit is highly homologous at both binding sites. The largest difference is the substitution of the foetal γ-subunit in our *T. californica* structure for the human adult ε-subunit^28^. This change of subunits results in different binding preference for the α-CTX: In *Torpedo*, α-GI and α-MI bind the α-δ interface with higher affinity than the α-γ interface as proposed from radioligand binding assays^16,29^. In adult mammalian (mouse) receptors, conotoxin α-MI also preferentially blocks the α-δ site^30^. Whether this preference holds for human adult muscle nAChR remains to be confirmed.

**Figure 3.**
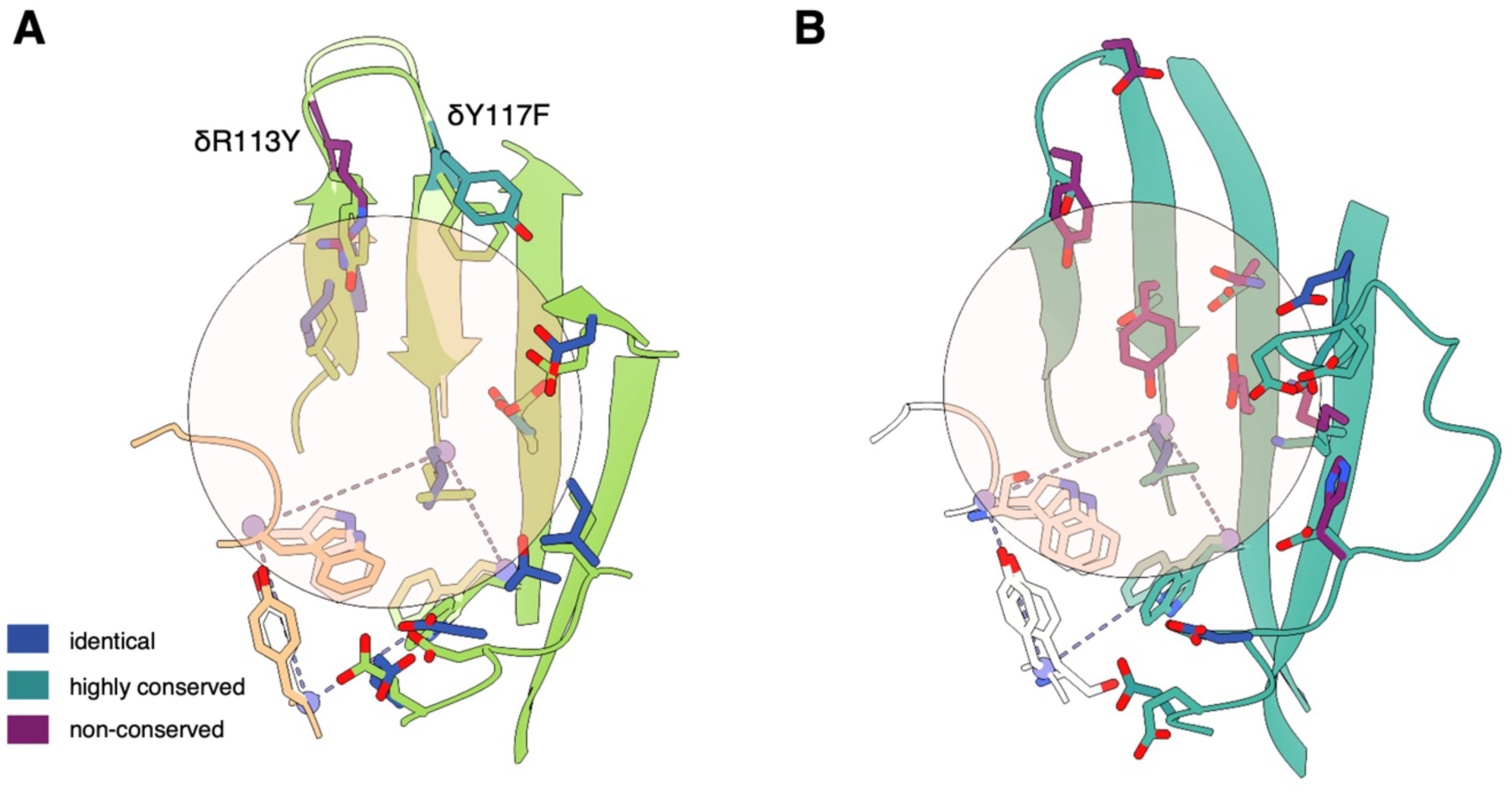
Comparison between *T. californica* and human nAChR α-CTX binding. (A) Schematic of the α-δ binding site and comparison between Torpedo and human nAChR sequences where interacting residues are identified by conservation between the two and the α-CTX binding is shown as a circle. (B) Schematic of the α-γ binding site and comparison between Torpedo and human nAChR sequences where interacting residues are identified by conservation between the two and the α-CTX binding is shown as a circle.

To better understand this potential switch in affinities and the binding of these toxins to the human muscle-type nAChR, we attempted to dock the α-CTXs used in this study at both sites of the apo adult-human muscle-type receptor (pdb: 9DMG)(Figure 3A). Docking at the α-δ interface yielded multiple structures with binding modes consistent with those experimentally determined in *T. californica.* Although many structures were consistent with experiment, only one of the consistent poses was highly ranked (inside the top ten best scores) and is shown in Figure 3A. However, docking was only successful at the α-δ interface as the α-ε binding site is by the residues between the β10 and Loop F regions. This Loop F region is shorter in the ε subunit than the δ and residue εN164 sits closer to the C-loop of the primary face than the closest residue in the δ subunit (δN169). This restricts binding site access preventing the conotoxins from docking at this interface (Figure S15) and is consistent with the selectivity for the human α-δ interface over the α-ε interface.

### α-CTX sequence diversity and conservation

Having established a model for α-CTX binding at muscle-type nAChR, we sought to determine if these binding features were conserved and could be used to infer information about receptor subtype specificity of α-CTXs. The sequences of all annotated α-CTXs with cysteine frameworks I and II (CC-C-C and CCC-C-C-C) were collected from the ConoServer database^5^ and aligned to produce a phylogenetic tree (Figure 4A).

**Figure 4.**
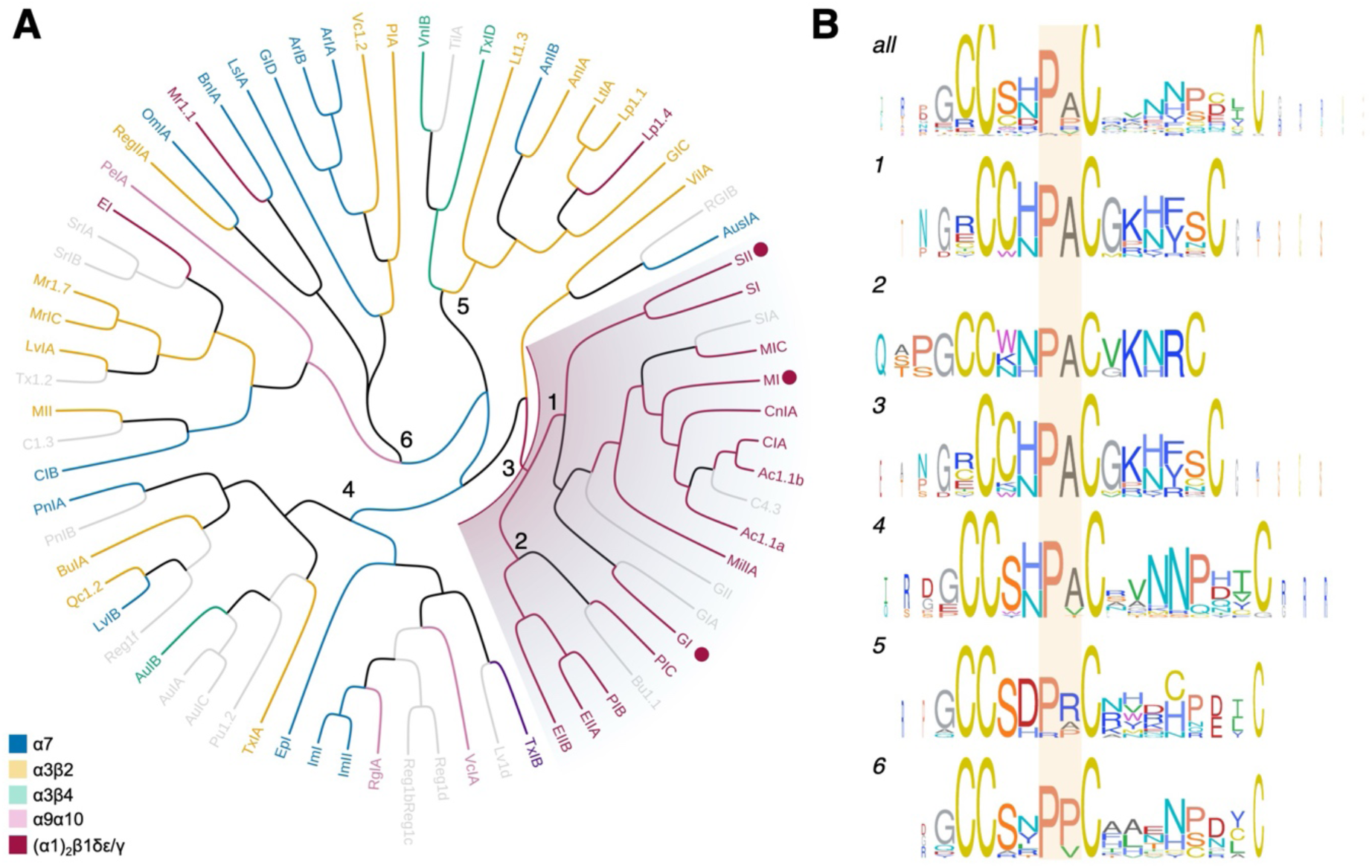
α-CTX sequence conservation groups toxins broadly along subtype specificity. (A) A phylogenetic tree of α-CTX sequences following alignment. Each toxin is colour coded to most potent receptor subtype as shown in the key, inset. Sequences from numbered nodes were subsequently examined for sequence conservation. α-CTXs in this study are identified by red circles within the shaded node. (B) Sequence alignments at the identified nodes were processed to produce logograms to examine sequence conservation. The Pro-0 and Ala+1 are highly conserved across much of α-CTX space. The 3/5 loop size is confined to muscle-type selective α-CTXs.

The resulting nodes broadly clustered the α-CTX sequences according to nAChR subtype specificity and enabled us to examine features present in α-CTXs targeting muscle-type nAChR that are not present in conotoxins targeting other subtypes. Within this set of sequences, we generally observe separation by the cysteine framework within the 3/5 α-CTXs shown in cluster 3. To highlight the most distinct features, we focused on narrow clusters of highly specific α-CTXs. The Pro-0/Ala+1 sequence is highly conserved and the defining feature in all clusters, aside from those α-CTXs active against the α3 containing nAChR in clusters 5 and 6 (Figure 4B). In α-conotoxins specific for muscle-type nAChRs, the residue preceding Pro-0 can be either histidine or asparagine, both capable of donating hydrogen bonds. In contrast, α-conotoxins targeting neuronal-type nAChRs are largely restricted to histidine at this position. Interestingly, cluster 5 contains aspartic acid in the -1 position and indicates a key difference in the binding mode of these toxins. Additionally, a glycine residue is favoured at position +3 in α-CTX specific for muscle type nAChRs that is not observed in α-CTX binding to other subtypes.

## DISCUSSION

Conotoxins form a vast reservoir of bioactive molecules that hold promise as therapeutics for a range of conditions, from pain management to neurological disorders^2^. A prerequisite for their rational development is a detailed molecular understanding of conotoxin/receptor interactions. These data improve our understanding of the exquisite specificity of conotoxins for their targets. In recent years, several structures of conotoxins in complex with their receptor(s) have been reported: the structures of voltage-gated calcium (Ca_v_) channels 2.1 and 2.2 in complex with ziconotide (ω-conotoxin MVIIA)^31^; the structure of the voltage-gated sodium channel Na_v_ 1.2 in complex with μ-conotoxin KIIIA^32^; and the noradrenaline transporter bound to analogues of the χ-conotoxin MrlA^33^. Despite this progress, structures for the largest class of pharmacologically classified conotoxins, α-CTXs have remained elusive.

We have determined three high-resolution structures of structurally diverse α-CTXs bound to muscle-type nAChR using cryo-EM. This has allowed us to describe their precise mode of binding and identify key structural moieties that drive potency and selectivity within muscle-type nAChRs (Figure 5). Each of the α-CTXs studied here bind within the ACh binding site and their binding to the receptor prevents inward movement of the C-loop, which precludes movement of the cys-loop and prevents pore opening^34^. α-CTXs engage in interactions across the binding interface between two subunits, with the α1-subunit acting as the principal binding site. This is in direct contrast to 3FTXs such as bungarotoxin^18^ and the scNTX^20^ for which binding is entirely reliant on the principal alpha subunit. As such the α-CTX binding mode itself contributes to selectivity across the nAChR family.

**Figure 5.**
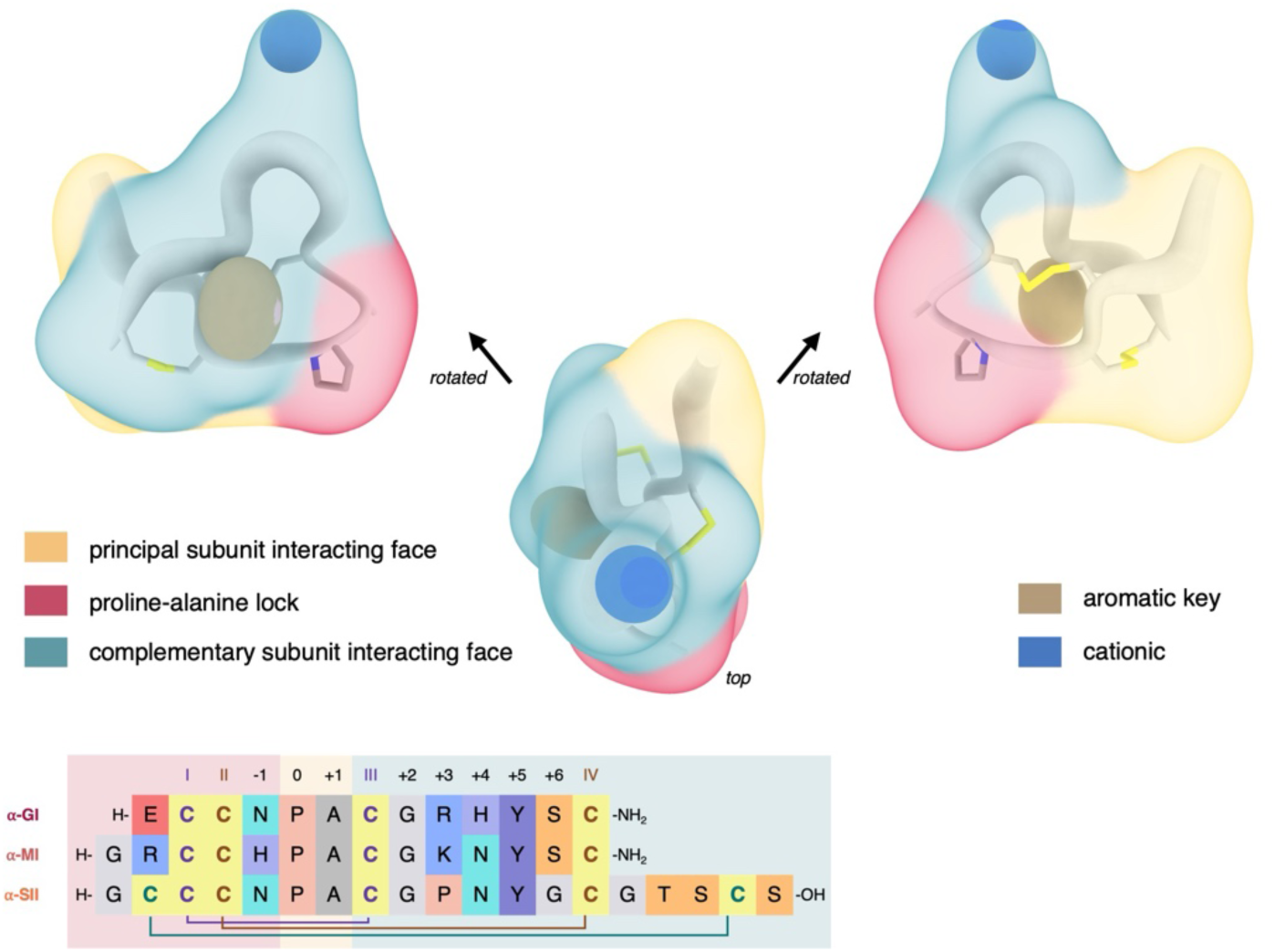
Generalised α-CTX (3/5) pharmacophore with conserved features that drive interaction at muscle-type nAChR. A model 3/5 α-CTX demonstrating surface interactions with the principal and complementary subunits of muscle-type nAChR. Conserved features are highlighted and sequence conservation across the 3/5 family is illustrated through cartoon ribbon thickness with conserved areas thinner and areas of higher sequence diversity thicker.

Each of the α-CTXs in this study binds at each nAChR site with similar topology. Both the N- and C-termini are solvent exposed whilst the highly conserved Pro-0 is buried firmly in the aromatic cage that forms the choline binding pocket. The previously published solution NMR ensembles of α-GI^35^ and α-SII^17^ are remarkably similar to the receptor-bound states we observe in our cryo-EM structures (Figure S12). Combined with the large number of hydrophobic interactions this conformational constraint suggests a low entropic cost of α-CTX binding, which contributes to their exquisite potency. The α-CTX II-IV disulfide bond is positioned near the vicinal disulfide bond of the C-loop. Intriguingly, we observe a connection between α-CTX CII and αC192 in the cryo-EM density. The importance of the CI-CIII disulfide bond for α-CTX bioactivity has been demonstrated across receptor subtypes through failed attempts to replace this particular disulfide bond with mimics that retain function, including triazole^36^, lactam^37^, thioether^38^ and diselenide bridges^39^. Interestingly, in all examples the remaining disulfide bond (CII-CIV) withstood replacement with a non-reducible surrogate functionality without affecting binding. While this observation in the map may be an artefact of disulfide reduction through electron-induced radiation damage, or reduction through exposure to reducing agent during purification, the possibility of the formation of an intermolecular disulfide bond is highly intriguing. It may also provide a rationale for the sustained paralysis following α-CTX envenomation on patients and could provide a significant contribution to α-CTX potency, and perhaps conotoxin function more broadly. More work is required to confirm this tentative mechanism.

The α-CTX numbering system centred on the central Pro-0 provides an easy way to delineate which part of the conotoxin binds to which receptor subunit: residues assigned negative integers engage the principal α1-subunit, while those assigned positive integers bind to the secondary subunit (δ and γ in our case). Our data allow us to propose generalized 3/5 α-CTX pharmacophores with conserved features: a central Pro-0/Ala+1 lock that is buried deep in the receptor interface, a face interacting with the principal subunit and a second face that interacts with the complementary subunit (Figure 5). This separation of subunit specific interactions in both structure and sequence provides a modular view of α-CTXs, with affinity for the α-subunit provided by residues *N-*terminal of Pro-0 and receptor subtype-specificity provided by the residues *C-*terminal of Pro-0.

In conclusion, our work defines a conserved α-CTX binding topology at muscle-type nAChRs and identifies key subunit-specific pharmacophores that drive potency and selectivity. These structural insights establish our molecular understanding of conotoxin recognition and provide a robust foundation for the rational design of future conotoxin-based therapeutics.

### Limitations of the study

While this work provides high-resolution structures of muscle-type nicotinic acetylcholine receptors (nAChRs) bound to diverse α-CTX, several limitations can be acknowledged. First, the structural data were obtained using *T. californica* nAChR as a surrogate for the human receptor. Although this model shares substantial sequence identity with the human adult receptor, differences in subunit composition, particularly the γ-to-ε substitution, may influence binding preferences and pharmacology. Consequently, extrapolation of these findings to human receptors should be approached with caution, and future studies using human nAChR structures will be required to validate these observations.

Second, the cryo-EM structures represent the receptor in a single closed resting state. While this provides valuable insight into antagonist binding, it does not capture the conformational dynamics associated with gating or desensitisation. Complementary approaches such as time-resolved cryo-EM or single-molecule electrophysiology could further elucidate these mechanisms.

Third, our pharmacological assays were performed in a model system and may not fully replicate the complexity of native neuromuscular junctions, where additional factors such as glycosylation, lipid composition, and accessory proteins could modulate toxin binding. Whilst this approach enables robust comparison of toxin activity, these assays reflect global change as opposed to single channel kinetics.

Together, these caveats highlight the need for integrative studies combining structural, functional, and physiological approaches to fully understand α-CTX selectivity and guide therapeutic development.

## RESOURCE AVAILABILITY

### Lead contact

Further information and requests for resources and reagents should be directed to and will be fulfilled by the lead contact, Andrew Jamieson.

### Materials availability

Requests for cell lines, plasmids and chemical reagents generated in this study should be directed to and will be fulfilled by Andrew Jamieson.

### Data and code availability

Cryo-EM maps and associated atomic models have been deposited in the EMDB and PDB respectively. α-GI bound nAChR (EMB-55142; 9SRL), α-MI bound nAChR (EMB-55143; 9RSM) and α-SII bound nAChR (EMB-55144; 9SRN).

## ACKNOWLEDGMENTS

We would like to thank Prof. David Beeson for supplying CN21ƔKO cells. We acknowledge the Scottish Centre for Macromolecular Imaging (SCMI) for access to cryo-EM instrumentation, funded by the MRC (MC_PC_17135, MC_UU_00034/7, MR/X011878/1) and SFC (H17007). We acknowledge Diamond for access and support of the cryo-EM facilities at the UK electron Bio-Imaging Centre (eBIC), proposal BI37630. M.A.M. and K.I.M.A. would like to acknowledge the EPSRC (EP/T517896/1 and EP/W524359/1) for sponsoring their Doctoral Scholarships. This work was funded by the US Defense Threat Reduction Agency (contract HDTRA12210001). The project or effort depicted was or is sponsored by the Department of Defense, Defense Threat Reduction Agency. The content of the information does not necessarily reflect the position or the policy of the federal government, and no official endorsement should be inferred.

## AUTHOR CONTRIBUTIONS

Conceptualization, A.G.J and J.K; Funding acquisition, J.W.E., J.G.F., J.K. and A.G.J.; Investigation, M.J.C., O.A.S., C.H., O.J.M., L.W., S.T., T.D.M, J.A.C.; Resources, O.A.S., N.W., M.A.M., J.W.E., J.K., A.G.J; Supervision, A.C.G., J.G.F., J.W.E, J.K., A.G.J.; Visualization, M.J.C.; Writing-original draft, M.J.C; Writing-review and editing, O.A.S., O.K.M., N.W., M.A.M. A.C.G, C.M.T., J.G.F., J.W.E, A.G.J and J.K;

## DECLARATION OF INTERESTS

The authors declare no competing interests.

## STAR★METHODS

### KEY RESOURCES TABLE

The items in the key resources table (KRT) must also be reported alongside the description of their use in the method details section. Literature cited within the KRT must be included in the references list. Please **do not edit the headings or add custom headings or subheadings** to the KRT. We highly recommend using <u>RRIDs</u> as the identifier for antibodies and model organisms in the KRT. To create the KRT, please use the template below or the <u>KRT webform</u>. See the more detailed <u>Word</u> <u>table template</u> document for examples of how to list items.

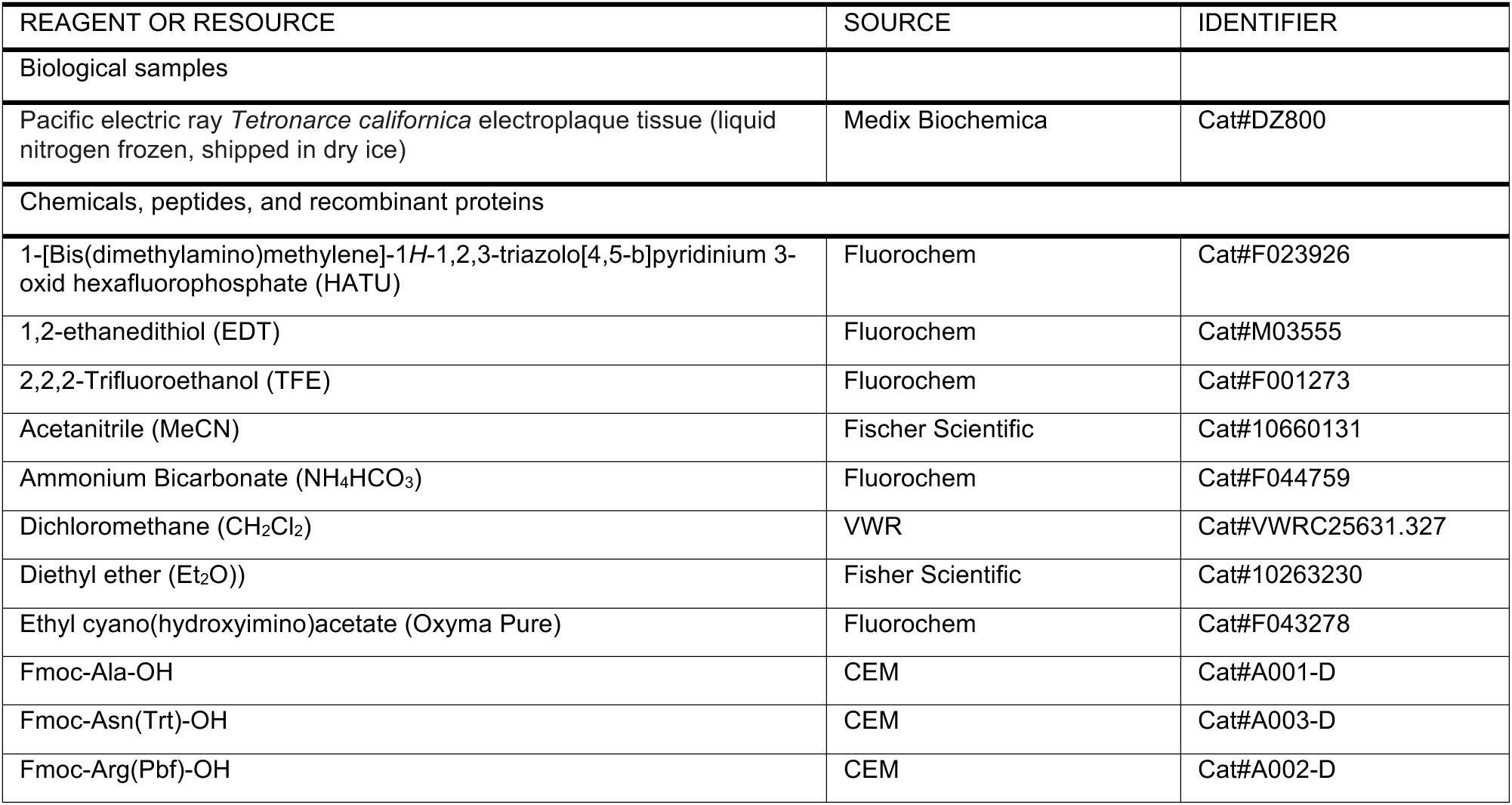
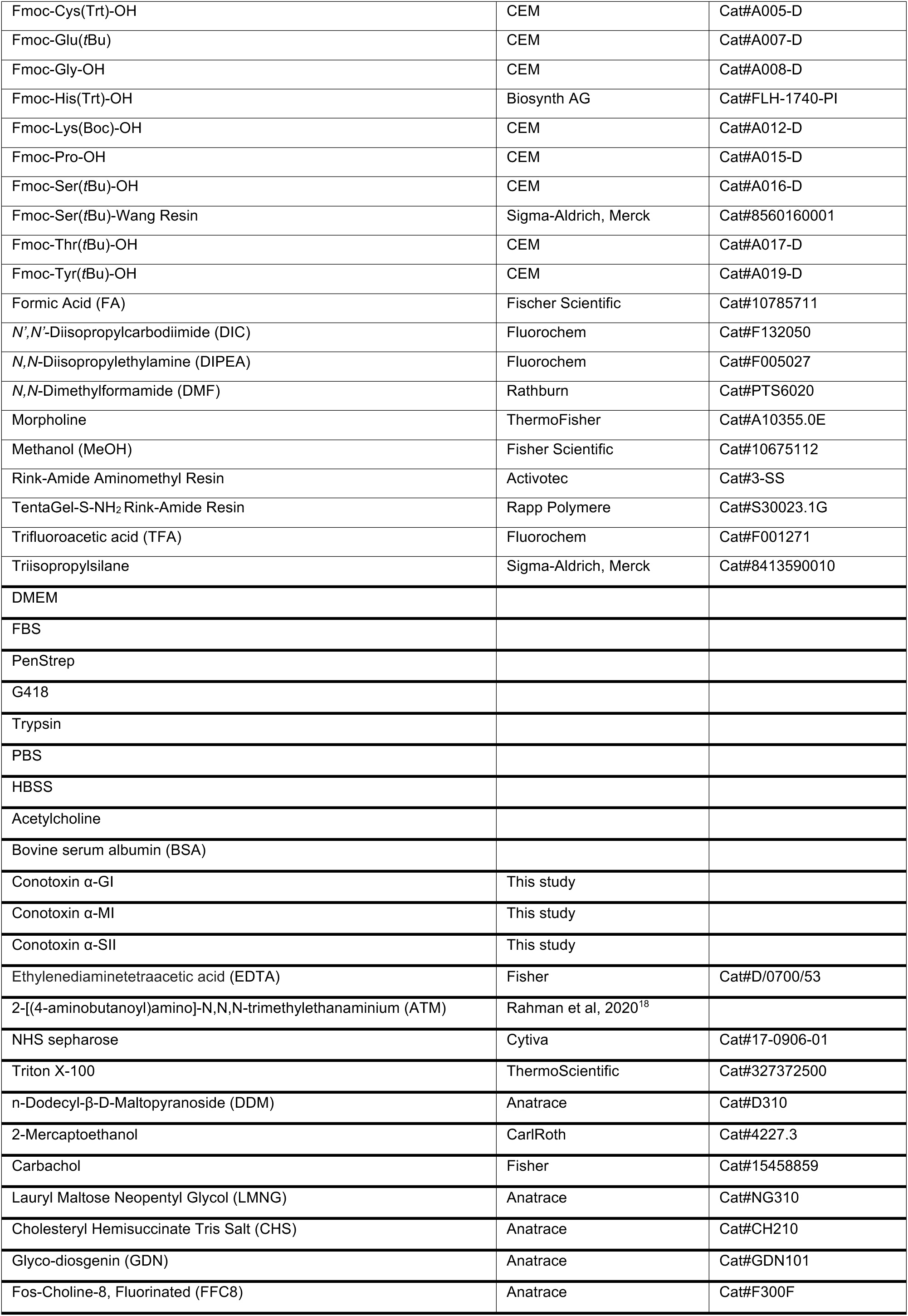
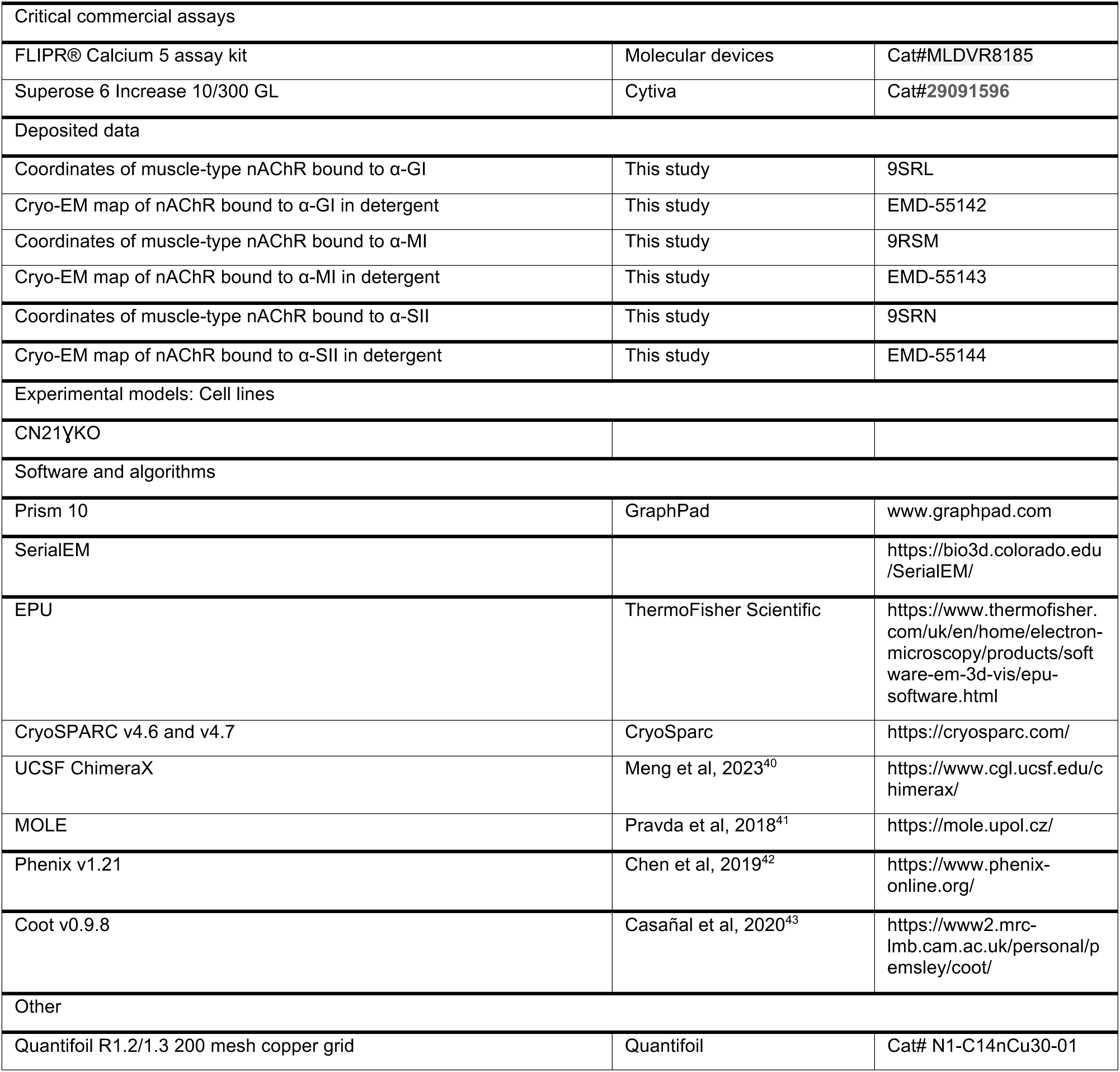

### METHOD DETAILS

#### General protocols for peptide synthesis

Peptide specific methods for all peptides synthesised for this work are in the supplementary methods. Solid phase peptide synthesis (SPPS) was used to produce all peptides described in this work.

#### Reagents and Instrumentation

All reagents were purchased from commercial sources and used without further purification unless stated. Standard Fmoc-protected amino acids were from CEM Corporation (Buckingham, UK) and Pepceuticals Ltd. (Leicester, UK), unless otherwise stated. Side-chain protecting groups of N^α^-Fmoc amino acids were; Fmoc-Asn(Trt)-OH (Trt = triphenylmethane), Fmoc-Asp(*t*Bu)-OH (*t*Bu = *tert*-butyl), Fmoc-Arg(Pbf)-OH (Pbf = 2,2,4,6,7-pentamethyldihydrobenzofuran-5-sulfonyl), Fmoc-Cys(Trt), Fmoc-Gln(Trt)-OH, Fmoc-Glu(*t*Bu), Fmoc-His(Trt)-OH, Fmoc-Lys(Boc)-OH (Boc = *tert*-butyloxycarbonyl), Fmoc-Ser(*t*Bu)-OH, Fmoc-Thr(*t*Bu)-OH and Fmoc-Tyr(*t*Bu)-OH.

Peptide grade *N,N*-Dimethylformamide (DMF) and reagent grade diethyl ether (Et_2_O) were from Rathburn Chemicals Ltd. (Walkerburn, Scotland). Triisopropylsilane (TIPS), was from Sigma-Aldrich/Merck Ltd. (Gillingham, UK) and *N,N*-diisopropylethylamine (DIPEA), 1,4-dioxane, hexafluorophosphate azabenzotriazole tetramethyl uronium (HATU), methanol-_d4_ (CD_3_OD), and 2,2,2-trifluoroethanol (TFE) were from Fluorochem (Glossop, UK). Morpholine was from Alfa Aesar Thermo Fischer Scientific Ltd. (Morecambe, UK). Dichloromethane (CH_2_Cl_2_) was from VWR International (Lutterworth, UK) and acetonitrile (MeCN), trifluoroacetic acid (TFA), and formic acid (FA) from Fisher Scientific. ChemMatrix^©^ Rink-Amide resin was obtained from Biotage (Hengoed, UK). TentaGel^®^ S Rink-Amide resin from Rapp Polymere GmbH (Tübingen, Germany) and aminomethyl Rink-Amide resin from Activotec (Toft, UK).

Analytical reverse-phase high-performance liquid chromatography (RP-HPLC) was performed on a Shimadzu RP-HPLC system with Shimadzu LC-20AT pumps, a Shimadzu SIL20A autosampler and a Shimadzu SPD-20A UV-vis detector using a Phenomenex Aeris Peptide XB-C18 column (100Å, 5 µm, 150 × 4.6 mm) (Macclesfield, UK). Compounds were eluted with linear gradients at column-dependent flow rates (1 mL/min for the Aeris), where buffer A = 0.1% TFA in Milli-Q^®^ (MQ) H_2_O and buffer B = 0.1% TFA in MeCN. Data are reported as column retention time (t_R_) in min. Crude peptides were purified by preparative RP-HPLC using an Agilent Technologies 1260 Infinity II Preparative LC System (monitoring at 214 nm and 280 nm) with a Phenomenex Gemini column 5 µm C18 column (100 Å, 250 x 21.2 mm). Peptides were eluted on linear gradients (10 mL/min) as determined by analytical RP-HPLC. Solvents employed were buffer; A = MQ H_2_O + 0.1% TFA, and buffer; B = MeCN + 0.1% TFA. Liquid chromatography-mass spectrometry (LCMS) was performed on a Thermo Scientific LCQ Fleet Ion Trap Mass Spectrometer using positive mode electrospray ionisation (ESI+). Where buffer A = 0.1% TFA in 95% MQ H_2_O/5% MeCN and buffer B = 0.1% TFA in 95% MeCN/5% MQ H_2_O, a linear gradient of 5%-95%B over 20 min with a flow rate of 1 mL/min was used, with a Phenomenex Aeris Peptide XB-C18 column (100 Å, 5 µm, 150 mm x 4.6 mm).

##### General Protocol 1 for the Automated Fmoc-SPPS

Peptides were synthesised batchwise as required on an Automated Biotage Initiator+ Alstra microwave synthesiser using preloaded Fmoc-Rink-Amide aminomethyl resin (0.53 mmol/g loading). Following initial resin swelling in DMF (4 mL) at room temperature (r.t.) for 30 min, and Fmoc removal, peptides were elongated in cycles of amino acid coupling followed by Fmoc removal. The coupling of Fmoc-protected amino acids (5 equiv., 0.2 M in DMF) was achieved by treatment with 1,3-diisopropylcarbodiimide (DIC, 5 equiv., 0.5 M in DMF) and ethyl cyano(hydroxyamino) acetate (Oxyma Pure, 5 molar equiv., 0.5 M in DMF) at r.t. for 60 min. Fmoc removal was achieved by treatment with 20% morpholine and 5% formic acid (*v/v*) in DMF (4 mL) at r.t (2 x 10 min). Following Fmoc removal and amino acid couplings, the resin was washed with DMF (4 x 4 mL), and after coupling (2 x 4 mL). Following final Fmoc removal, the resin was removed and manually washed with DMF (3 x 5 mL), MeOH (3 x 5 mL), CH_2_Cl_2_ (3 x 5 mL), and dried under vacuum.

##### General Protocol 2 for the Manual Fmoc-SPPS

Preloaded Fmoc-Ser(*t*Bu)-Wang Resin (0.667 g, 0.2 mmol, 0.3 mmol/g) was swollen in CH_2_Cl_2_ at r.t. for 10 min followed by DMF at r.t. for 10 min. For Fmoc removal, the peptidyl resin was twice treated with a solution of 20% morpholine and 5% formic acid (*v/v*) in DMF at r.t for 10 min. Fmoc amino acids (0.8 mmol, 4.0 equiv) were employed to elongate the peptide alongside HATU (289 mg, 0.76 mmol, 3.8 equiv.) and DIPEA (348 μL, 2 mmol, 10 equiv.) in DMF at r.t. for 60 min. Following Fmoc removal and amino acid couplings, the peptidyl resin was washed with DMF (3 x 5 mL), MeOH (3 x 5 mL), CH_2_Cl_2_ (3 x 5 mL), and again with DMF (3 x 5 mL). Upon complete elongation of the peptide, the resin was washed with DMF (3 x 5 mL), MeOH (3 x 5 mL), CH_2_Cl_2_ (3 x 5 mL), and dried under vacuum.

##### General Protocol 3 for Resin Cleavage and Global Deprotection

The resin-bound peptide was treated with a cleavage cocktail of TFA/H_2_O/TIPS (95/2.5/2.5, *v/v/v*, 10 mL per 0.1 mmol resin) and agitated (1-2 h, r.t.). The cleavage solution was separated from the resin and its volume reduced under a flow of nitrogen. Ice cold Et_2_O (15 mL per 0.1 mmol resin) was used to precipitate the peptide. The precipitate was isolated by centrifugation, washed with ice cold Et_2_O, dissolved in MeCN/MQ H_2_O (2:8, *v/v*) and analysed by RP-HPLC and LCMS. Following determination of the desired peptidyl products, the solutions were frozen and lyophilised.

##### General Protocol 4 for Solution Phase Oxidation

The lyophilised peptide was solubilised in a solution of either MeCN/MQ H2O (1:1, v/v) or TFE/MQ H2O (1:1, v/v) buffered to pH ∼9.5 by 0.1 M NH_4_HCO_3_. The respective linear conotoxin was added over 2 h with mixing at r.t. overnight. Complete oxidation was confirmed by analytical RP-HPLC and LCMS.

#### ACh calcium response assays

Calcium response assays were carried out as previously established^15,36^. CN21ɣKO cells expressing only the adult isoform of muscle-type nAChR were grown in DMEM supplemented with 10% FBS, Glutamax and 1% P/S in TC treated dishes at 37°C, 5% CO_2_. For assay, cells were seeded at 20,000 or 40,000 per well in black, clear bottom 96 well plates and incubated for 48 or 24 hours prior to assay respectively. On the day of assay, media was removed cells were loaded with calcium FLPR dye (Molecular Devices) resuspended in 20 mM HEPES pH 7.5, 1x HBSS and diluted 2-fold with DMEM before the addition of 20 µM atropine to prevent muscarinic receptor mediated response. Cells were then incubated at 37°C for 25 min prior to the addition of α-CTX which were then incubated for a further 5 min. Plates were then placed in a prewarmed FlexStation 3 and fluorescence was measured at an excitation and emission wavelength of 485 and 525 nm respectively, with a cut-off at 515 over 60 s. 150 µM ACh, was added after 16 s and the resulting fluorescence upon calcium release was measured. The average of the 15 s baseline was subtracted from the maximum fluorescence and plotted against dose to form a dose response curve from which an IC_50_ could be calculated using GraphPad Prism10 and a four-parameter fit. Concentrations were run in quadruplicate on each plate, and each experiment was run independently at least four times.

#### Protein preparation and purification

*T. californica* nAChR was purified following established protocols^18,19^. Briefly, frozen electric organ tissue (Medix Biochemica, Espoo, Finland) was thawed before being resuspended in 20 mM NaH_2_PO_4_ pH 7.4, 400 mM NaCl, and 0.5 mg/ml *N*-ethylmaleimide and homogenised in a blender. Membranes were isolated by centrifugation at 200,000 g for 25 min, resuspended in 20 mM Tris pH 11, 80 mM NaCl, 1 mM EDTA, 20% sucrose and incubated for 30 min on ice. Membranes were harvested and homogenised in 20 mM NaH_2_PO_4_ pH 7.4, 100 mM NaCl thrice undergoing multiple washes. Membranes were snap frozen in liquid nitrogen and stored at -80 °C.

Membranes were resuspended in 20 mM NaH_2_PO_4_ pH 7.4, 80 mM NaCl and solubilised in 1.5% Triton X-100 for 1 h before being spun at 200,000 g for 30 min to remove insoluble material. Solubilised material was incubated with immobilised ligand affinity resin (ATM-sepharose) for 1 h before being packed and washed in 20 mM TIS pH 8, 100 mM NaCl, 1 mM ethylenediaminetetraacetic acid (EDTA), 0.05% *n-*dodecyl-*b*-d-maltopyranoside/cholesteryl hemisuccinate (DDM:CHS) – CHS was omitted from purification of protein used in the structure of α-SII. Protein was eluted in the same buffer containing 100 mM carbachol and 10 mM 2-mercaptoethanol. Protein-containing fractions were concentrated and desalted to remove the carbachol and 2-mercaptoethanol.

Protein for cryo-EM was then incubated with 0.1% lauryl maltose neopentyl glycol (LMNG):CHS on ice for 1 hr before size-exclusion chromatography (SEC) on a Superose 6 10/30 in 20 mM 4-(2-hydroxyethyl)-1-piperazineethanesulfonic acid (HEPES) pH 7.5, 100 mM NaCl, 0.1 mM EDTA, 0.00075% LMNG, 0.00015% CHS and 0.00015% GDN (glycol-diosgenin).

#### Cryo-EM preparation and data acquisition

SEC purified nAChR was concentrated to 5-10 µM and kept on ice. α-CTX were dissolved to 10 mM in SEC buffer before being added to purified receptor at 3-fold excess and incubated on ice for 30 min. The complex was concentrated to *ca.* 6 mg/mL (20-30 µM) for freezing and 0.3 mM FFC8 was added to prevent preferred orientation of particles in the ice. 3 µl of protein was added to Quantifoil 1.2/1.3 300 Cu mesh grids that had been glow discharged in air for 30 s at 35 mA. Samples were plunge-frozen in liquid ethane using a vitrobot with blot force 3 for 3 s at 8°C and 95% humidity. Frozen grids were stored in liquid nitrogen before screening and data collection.

Grids were screened on a JEOL F200 at SCMI (Glasgow, UK). Those demonstrating thin ice were used for high-resolution data collection on a JEOL CRYOARM300 (SCMI, Glasgow, UK) or eBIC (Diamond, Oxford, UK). Data collection statistics appear in Table S1.

#### Data processing and model refinement

Data were processed in cryoSPARC v4.6.2^44^. Individual workflows are detailed in Figures S2-S4 and model statistics in Table 1. Each dataset was processed similarly beginning with Patch Motion Correction and Patch CTF Estimation. Particles were picked with templates generated from screened grids on a JEOL F200 and extracted 4x binned prior to two rounds of 2D classification. Junk particles were discarded, and *ab initio* models were produced (from the initially discarded junk particles to act as sinks and the particles following classification). Heterorefinement of all particles in the 2D classified stack and those fitting the sole remaining good class were re-extracted 2x binned. Particles were CTF refined and subjected to a masked 3D classification to remove the detergent micelle which yielded multiple maps. The best group of particles were then extracted un-binned following reference-based motion correction and subjected to non-uniform (NU) refinement, yielding a final volume which was sharpened. Resolutions were determined according to gold-standard Fourier shell correlation (GS-FSC) before further examination of local resolution with a cut-off of 0.143.

The model 7smm (apo *T. californica* nAChR)^45^ was fitted in the map and refined using Phenix^42^ followed by manual building and the addition of the conotoxins, lipids, glycosylations and waters in Coot^43^ followed by further real-space refinement in Phenix. Final models were deposited in the Protein Data Bank (PDB) and validated. Protein interface area was calculated using ePISA^46^and pore analysis using MOLEonline^41^. Images were created in ChimeraX^47^ and interactions analysed in arpeggio^48^.

#### Sequence alignments

Sequence alignment was carried out using Clustal Omega^49^. Logograms to display conservation where produced using Weblogo^50^. TreeViewer was used to generate a tree from the sequence alignments^51^.

#### Molecular docking

The α-CTXs resolved in the structures presented here were docked to the extracellular region of the human adult nAChR taken from the PDB (ID: 9DMG). The missing residues in the β-subunit were modelled in using Modeller^52^. All docking was performed using the HADDOCK 2.4 webserver^53^. Active residues were defined from the list of interacting residues between the α-CTX and the receptor excluding water mediated interactions (Table S3 and S4). Three repeats were run and the default settings for protein-peptide docking were used with the addition of refining the final structures with 1 ns of molecular dynamics simulation in explicit solvent as per the HADDOCK final refinement protocol. The sample size was increased by setting the number of structures allowed in each stage of the docking process to their maximum (10,000 in it0 and 1,000 in it1). The final 200 structures from each repeat were then analysed by docking score and RMSD between α-CTX and the experimental α-CTXs bound receptor structure, with the best scoring structure with the lowest RMSD being presented in this work.

#### Water modelling

The computational water placement was performed using grand canonical Monte Carlo (GCMC) simulation using both instantaneous and nonequilibrium move proposals^54^ as implemented in the *grand* Python library^55^. To setup the systems, all glycans were removed from the cryo-EM structure and the subunits were cut such that only the extracellular region was simulated. For the alpha subunits, the proteins were cut from residue Pro211 onwards, the beta subunit was cut from residue Leu218 onwards, the delta subunit from residue Leu226 onwards and the gamma subunit from residue Leu220 onwards. In each case, an NME cap was added to ensure the termini remained neutral. The system was protonated at pH 7.4, solvated and parameterized in accordance with the AMBER 14ffSB forcefield^56^ for the protein and ions and the TIP3P forcefield^57^ for the water molecules. Sodium and chloride ions were added to both ensure the system was neutral and achieve an ion concentration of 0.15 M.

The simulations were performed in OpenMM 8.0.0^58^. Equilibration was performed over a series of steps combining both GCMC moves and molecular dynamics (MD) simulation. An initial 10,000 GCMC moves were followed by 100 cycles of 1,000 GCMC moves and 10 fs MD in the NVT ensemble. 5 ns of MD in the NPT ensemble was then used to ensure the system volume was equilibrated correctly. Finally, 500 cycles of 1 ps NVT simulation and 200 GCMC moves completed the equilibration. The production simulations were performed using nonequilibrium GCMC (GCNCMC) moves – each simulation consisting of approximately 15,000 moves with 5 ps MD sampling between moves. To analyse the water placement within the simulations, the trajectories were decorrelated such that every 10^th^ frame was retained and a hierarchal average-linkage clustering algorithm, as implemented in SciPy^59^, was used to determine the locations and occupancies of each cluster.

The GCMC moves were only performed within a spherical region within each system. For each binding site, the sphere is centered on the midpoint of the C-alpha atoms of two selected residues. In all cases the sphere radius was 14 Å. Methods table 1 shows the atoms chosen to define the sphere for each structure.

**Methods table 1.** The residues chosen to define the spherical region within which the water sampling was enhanced. In each case the position of the C-alpha atom of the residues indicated was used. For the α-GI and α-SII structures, the same residues were used at both binding sites.

| Structure | Atom 1 | Atom 2 |
| --- | --- | --- |
| α-MI $\delta$ | Cys4 | Cys8 |
| α-MI $\gamma$ | Cys3 | Tyr12 |
| α-GI δ-γ | Asn4 | Tyr11 |
| α-SII δ-γ | Asn5 | Cys8 |

Harmonic positional restraints were used to prevent movement of the α-GI conotoxin in the α-δ binding site, as was observed in initial simulations. These restraints were applied to all C-alpha atoms in both the α-CTX and the receptor and a force constant of 100 kJ nm^-2^ was used. No restraints were applied to the α-γ binding site.

## SUPPLEMENTAL INFORMATION

Document S1. Figures S1 – S25, Tables S1 – S5 and additional peptide methods.

## Notes

### Competing Interest Statement

The authors have declared no competing interest.

