## Supplemental Information for "Structures of muscle-type nicotinic acetylcholine receptor with α-conotoxins reveal determinants of receptor-subtype specificity"

*Supplemental Document S1*

Tables S1-S4
Figures S1-S15

*Additional peptide methods*
Table S5
Conotoxin Syntheses
Figures S16-S25

**Table S1. Table of cryo-EM data collections and model building statistics.**

| | **nAChR α-GI**<br>PDB 9SRL<br>EMD-55142 | **nAChR α-MI**<br>PDB 9RSM<br>EMD-55143 | **nAChR α-SII**<br>PDB 9SRN<br>EMD-55144 |
|---|---|---|---|
| **Data collection** | | | |
| EM facility | SCMI | eBIC | eBIC |
| Magnification | 60,000x | 165,000x | 165,000x |
| Voltage (kV) | 300 | 300 | 300 |
| Number of frames | 45 | 60 | 60 |
| Electron exposure ($e^-/Å^2$) | 45.0 | 60.0 | 46.3 |
| Target defocus (μm) | -0.5 *to* -2.0 | -0.5 *to* -2.0 | -0.5 *to* -2.0 |
| Pixel size (Å) | 0.7832 | 0.504 | 0.504 |
| Micrographs | 25,190 | 17,850 | 23,077 |
| **Reconstruction** | | | |
| Initial particles | 1,336,946 | 2,299,548 | 935,386 |
| Final particles | 303,864 | 177,883 | 358,667 |
| Symmetry | C1 | C1 | C1 |
| Box size (pixels) | 512 | 512 | 512 |
| GS-FSC threshold | 0.143 | 0.143 | 0.143 |
| Map resolution (Å) | 3.00 | 2.21 | 2.57 |
| Map sharpening B factor ($Å^2$) | 116.9 | 59.1 | 102.2 |
| **Refinement** | | | |
| Overall Q-score | 0.70 | 0.79 | 0.68 |
| Molprobity score | 1.45 | 1.26 | 1.27 |
| Clash score | 4.42 | 3.07 | 2.90 |
| Poor rotamers (%) | 0.00 | 0.00 | 0.53 |
| RMSD values | | | |
| Bond lengths (Å) | 0.006 | 0.006 | 0.005 |
| Bond angles (˚) | 0.568 | 0.627 | 0.706 |
| Ramachandran | | | |
| Favored (%) | 96.43 | 97.12 | 96.82 |
| Allowed (%) | 3.57 | 2.88 | 3.17 |
| B-factors ($Å^2$) | | | |
| Protein | 50.39 | 46.35 | 55.04 |
| Ligands | 76.73 | 69.99 | 51.36 |
| Model composition | | | |
| Non-hydrogen atoms | 18,206 | 18,543 | 12,229 |
| Protein residues | 2,070 | 2,073 | 1,416 |
| Ligands | | | |
| Waters | 450 | 780 | 191 |

**Table S2. Measured surface area and interfaces of α-CTX at each of the ACh binding sites. A comparison to well-studied toxin α-bungarotoxin is included.**

| | Surface area | α-δ interface | δ interface | Total interface | Surface area | α-γ interface | γ interface | Total interface |
|---|---|---|---|---|---|---|---|---|
| **α-GI** | 1312.5 | 375.7 | 427.62 | 803.32 (61.2%) | 1328.9 | 372.1 | 447.82 | 819.92 (61.7%) |
| **α-MI** | 1397.2 | 401.52 | 515.81 | 917.33 (65.7%) | 1441.2 | 402.55 | 568.02 | 970.57 (67.3%) |
| **α-SII** | 1590.7 | 376.2 | 526.78 | 902.98 (56.8%) | 1568.9 | 387.2 | 457.47 | 844.67 (53.8%) |
| **α-BTX** | 5186.5 | 845.79 | 685.36 | 1531.15 (29.5%) | 5190.1 | 783.31 | 716.44 | 1499.75 (28.9%) |

Interface areas are measured in $Å^2$. α-BTX taken from PDB 6uwz.

**Table S3. Molecular interactions between α-CTX and the α-δ binding site.**

| | | α-GI | α-MI | α-SII |
|---|---|---|---|---|
| *Principal face* | | | | |
| α | *N'* | Y190 | Y190 | Y190 |
| | I | Y190, C192 | Y190, C192 | Y190, C192 |
| | II | Y190, K145* | Y190 | Y190 |
| | -1 | I148*, Y151*, Y198 | Y93, Y198 | I148*, Y190*, Y193* |
| | 0 | Y93, W149 | Y93, W149 | Y93, W149 |
| | +1 | Y198* | Y198 | Y198* |
| | III | Y198 | Y198 | Y198 |
| | +2 | | | |
| | +3 | | | |
| | +4 | | | |
| | + | | | |
| | +6 | | | |
| | IV | | | |
| | *C'* | | | |
| *Complementary face* | | | | |
| δ | *N'* | | D180 | D165 |
| | I | | | |
| | II | | | |
| | -1 | | | |
| | 0 | W57, L121, N109* | W57, L121, N109* | W57, L121, N109* |
| | +1 | | | |
| | III | R113* | R113* | |
| | +2 | D59* | D59* | D59* |
| | +3 | S76*, D117*, D165 | D59*, R113, Y117 | D59*, R113 |
| | +4 | | | |
| | +5 | S36*, T38, W57, L121 | T38, W57, L121, D165 | T38, W57, L121 |
| | +6 | D165, | D165 | |
| | IV | I178, D180 | I178, D180 | |
| | *C'* | D176* | | |

*Water mediated

**Table S4. Molecular interactions between α-CTX and the α-ɣ binding site.**

| | | α-GI | α-MI | α-SII |
|---|---|---|---|---|
| *Principal face* | | | | |
| α | *N'* | Y190 | Y190 | |
| | I | Y190, C192 | Y190, C192 | |
| | II | Y190 | Y190 | |
| | -1 | Y93, Y198 | Y93, Y198, L199 | |
| | 0 | Y93, N107*, W149, W149* | Y93, N107*, W149, | |
| | +1 | Y151, Y198, Y198* | W149, Y151, Y198 | |
| | III | Y190, Y198 | Y190, Y198 | |
| | +2 | | | |
| | +3 | | | |
| | +4 | | | |
| | + | | | |
| | +6 | | | |
| | IV | | | |
| | *C'* | | | |
| *Complementary face* | | | | |
| γ | *N'* | | D174, E176 | |
| | I | | | |
| | II | | | |
| | -1 | | | |
| | 0 | W55, L119 | K34*, W55, L119, Y117* | |
| | +1 | | | |
| | III | Y111* | Y111* | |
| | +2 | | Y117 | |
| | +3 | E57*, Y111, Y111*, D113 | Q59*, Y111, S115, Y117 | |
| | +4 | | | |
| | +5 | W55, E57*, L119, | K34, T36, W55, L119 | |
| | +6 | | E164 | |
| | IV | D174 | D174 | |
| | *C'* | T36 | H172 | |

*Water mediated

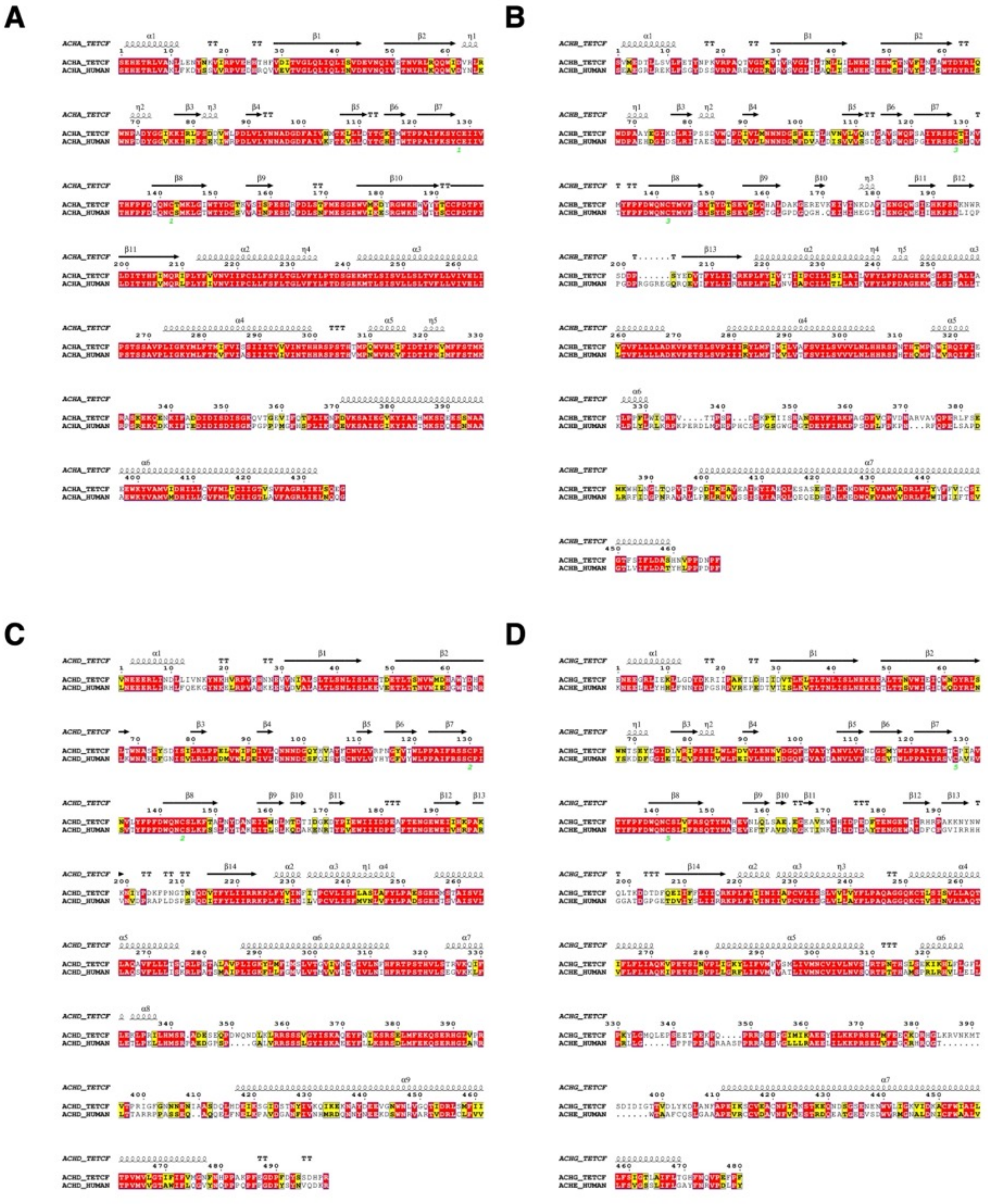

**Figure S1. Sequence alignments for human and *T. californica* nAChR subunits.**
(A) Alpha subunit
(B) Beta subunit
(C) Delta subunit
(D) Alignment between Torpedo gamma subunit and adult human epsilon

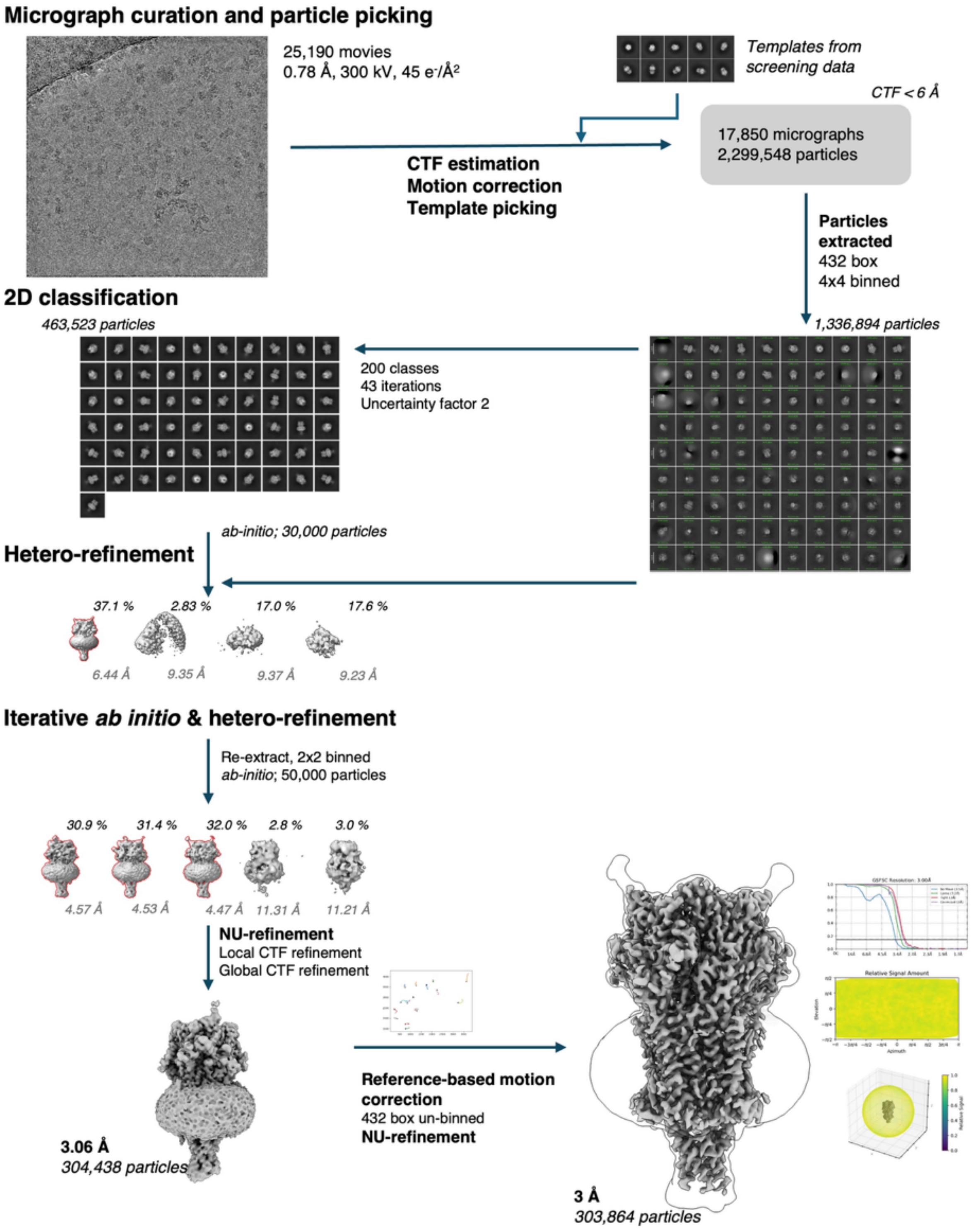

**Figure S2. Cryo-EM data processing flowchart with orientation statistics and GS-FSC curve for the final α-GI bound nAChR structure.**
All processing was carried out in cryoSPARC v4.6.2.

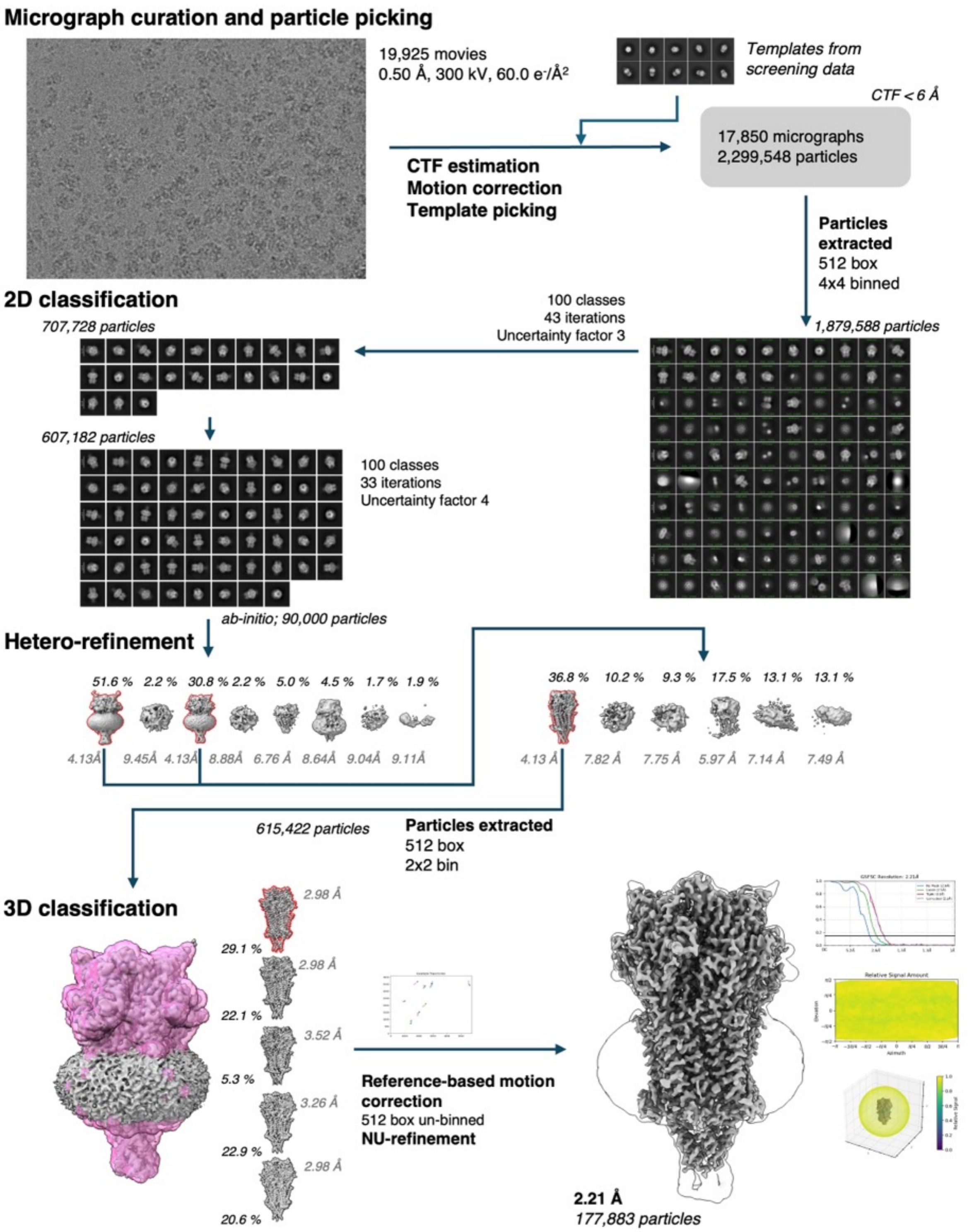

**Figure S3. Cryo-EM data processing flowchart with orientation statistics and GS-FSC curve for the final α-MI bound nAChR structure.**
All processing was carried out in cryoSPARC v4.7.

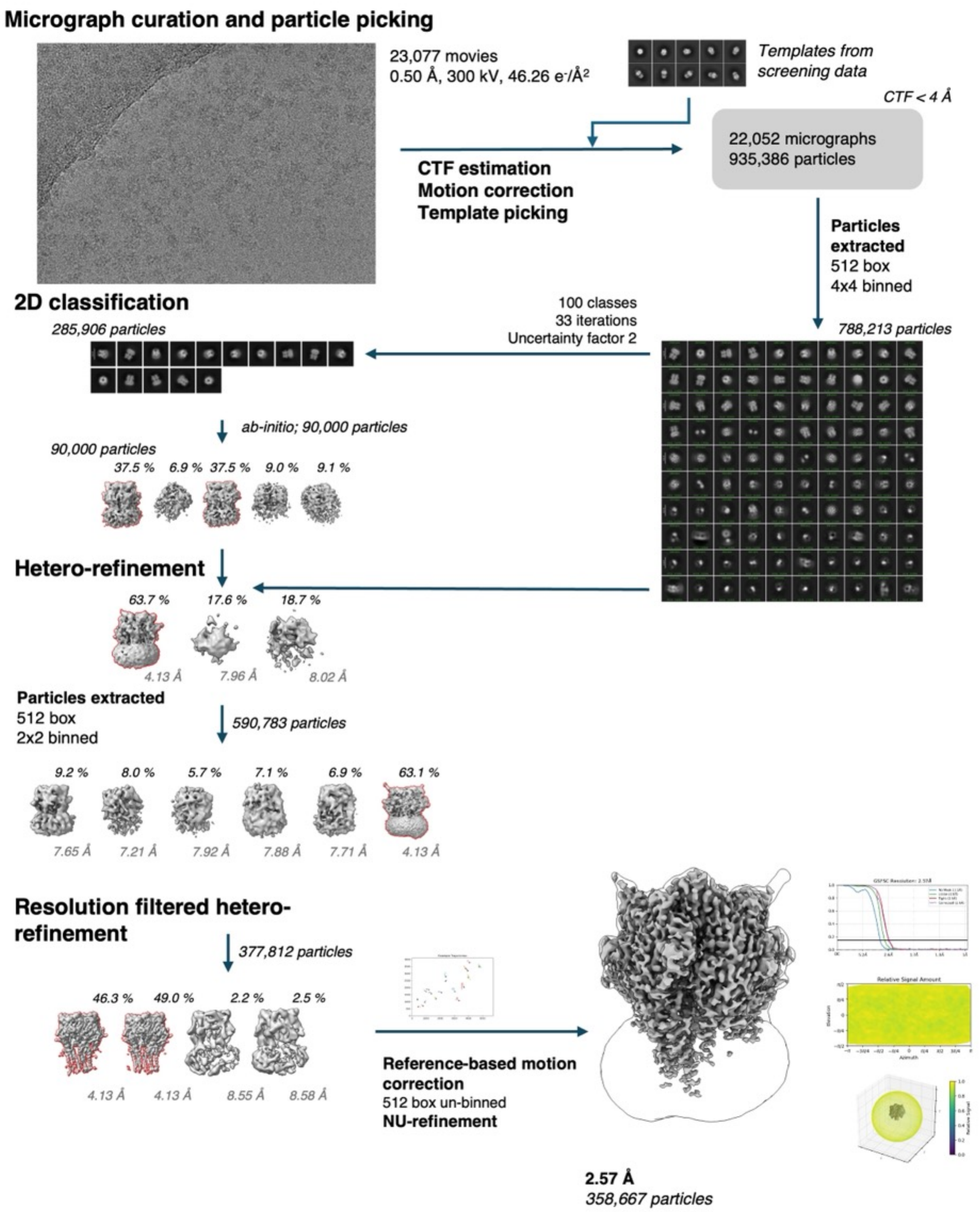

**Figure S4. Cryo-EM data processing flowchart with orientation statistics and GS-FSC curve for the final α-SII bound nAChR structure.**
All processing was carried out in cryoSPARC v4.7.

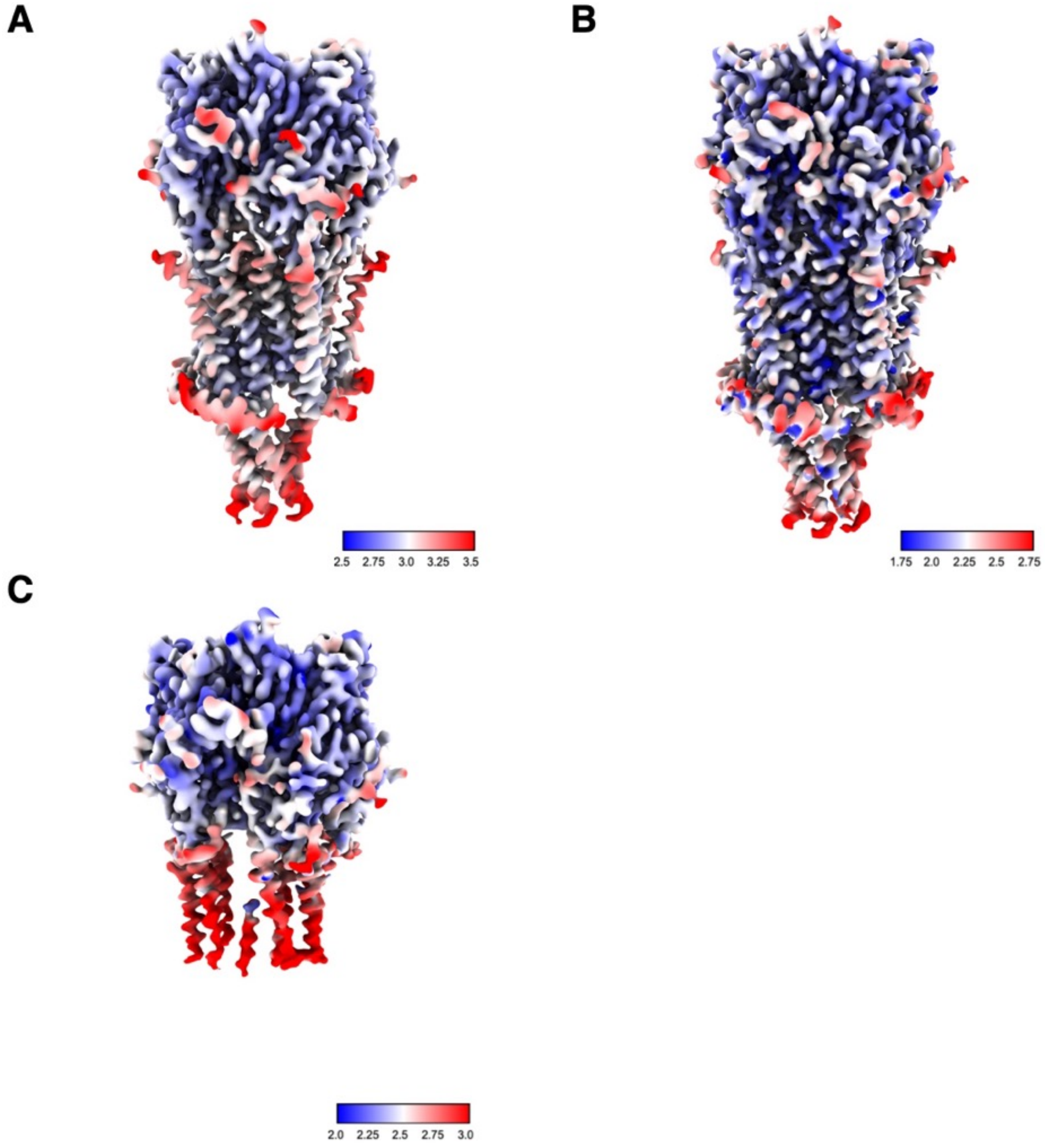

**Figure S5. Local resolution maps for each of α-CTX bound nAChR structures.**
Each of the conotoxin-bound nAChR structure maps colored by local resolution calculated in cryoSPARC with a GS-FSC cut-off of 0.143. (A) α-GI. (B) α-MI. (C) α-SII. Color legends show resolution in Å.

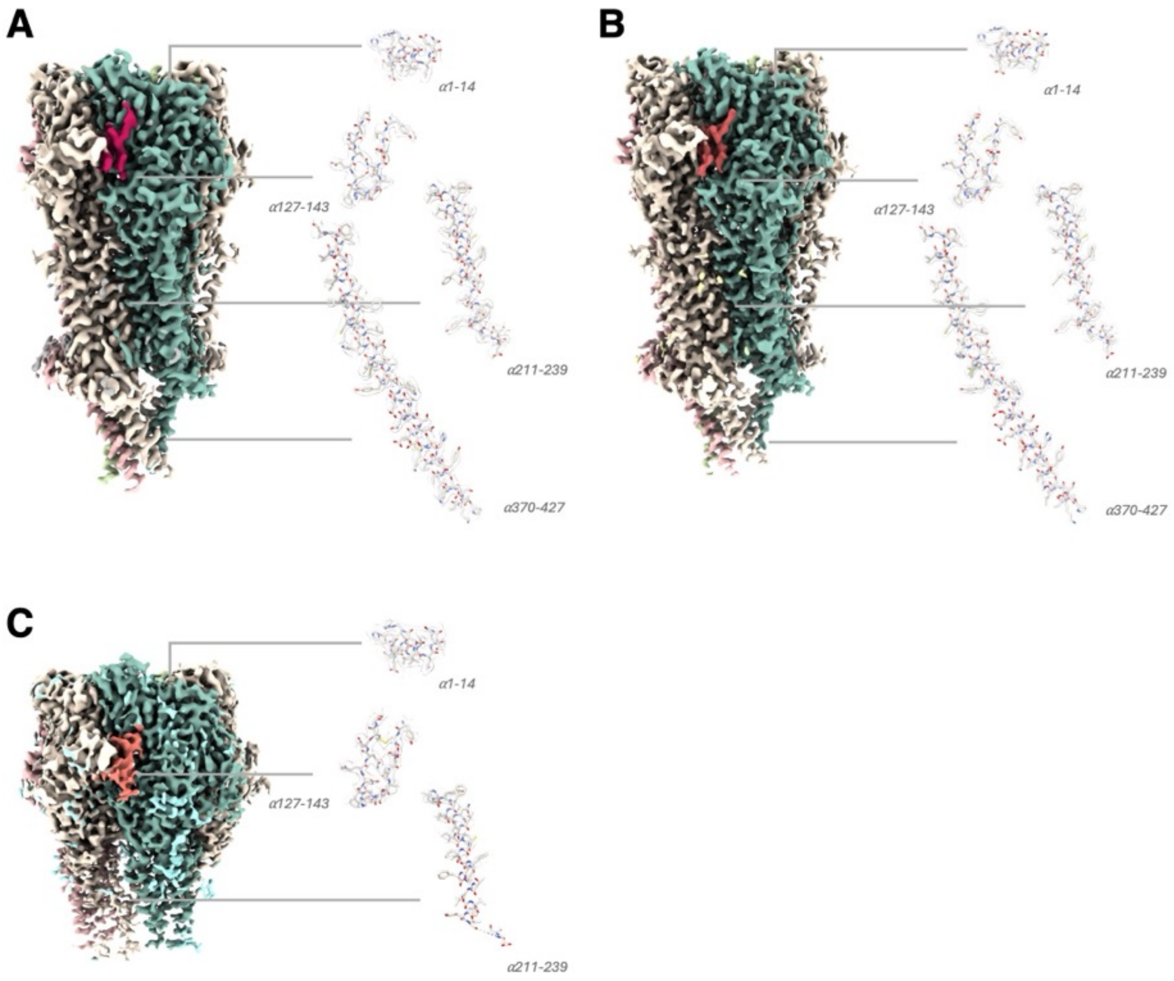

**Figure S6. Sharpened cryo-EM density maps contoured and surrounding areas of nAChR.**
(A) α-GI. (B) α-MI. (C) α-SII.

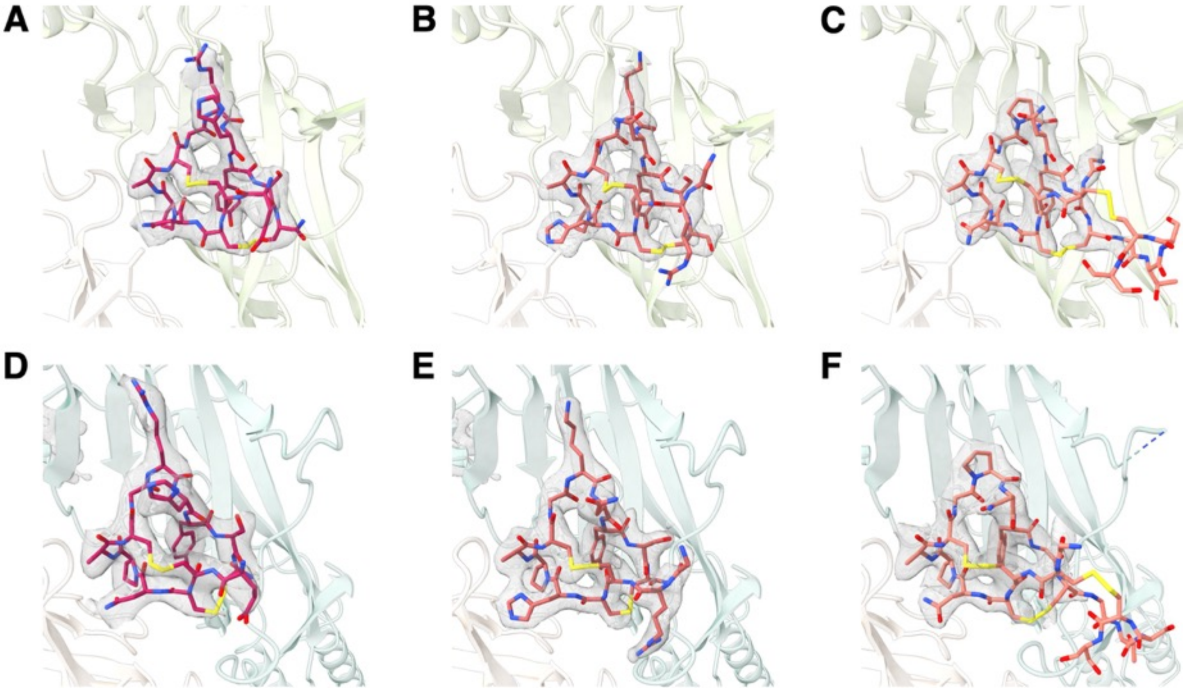

**Figure S7. Density maps of each α-CTX at both binding sites in muscle-type nAChR.** Sharpened cryo-EM density maps contoured and surrounding each α-CTX at α-δ and α-ɣ sites respectively (A and D) α-GI. (B and E) α-MI. (C and F) α-SII. The additional C'-terminal loop of α-SII was not visible in the sharpened map although density confirmed it as present in the unsharpened map from refinement.

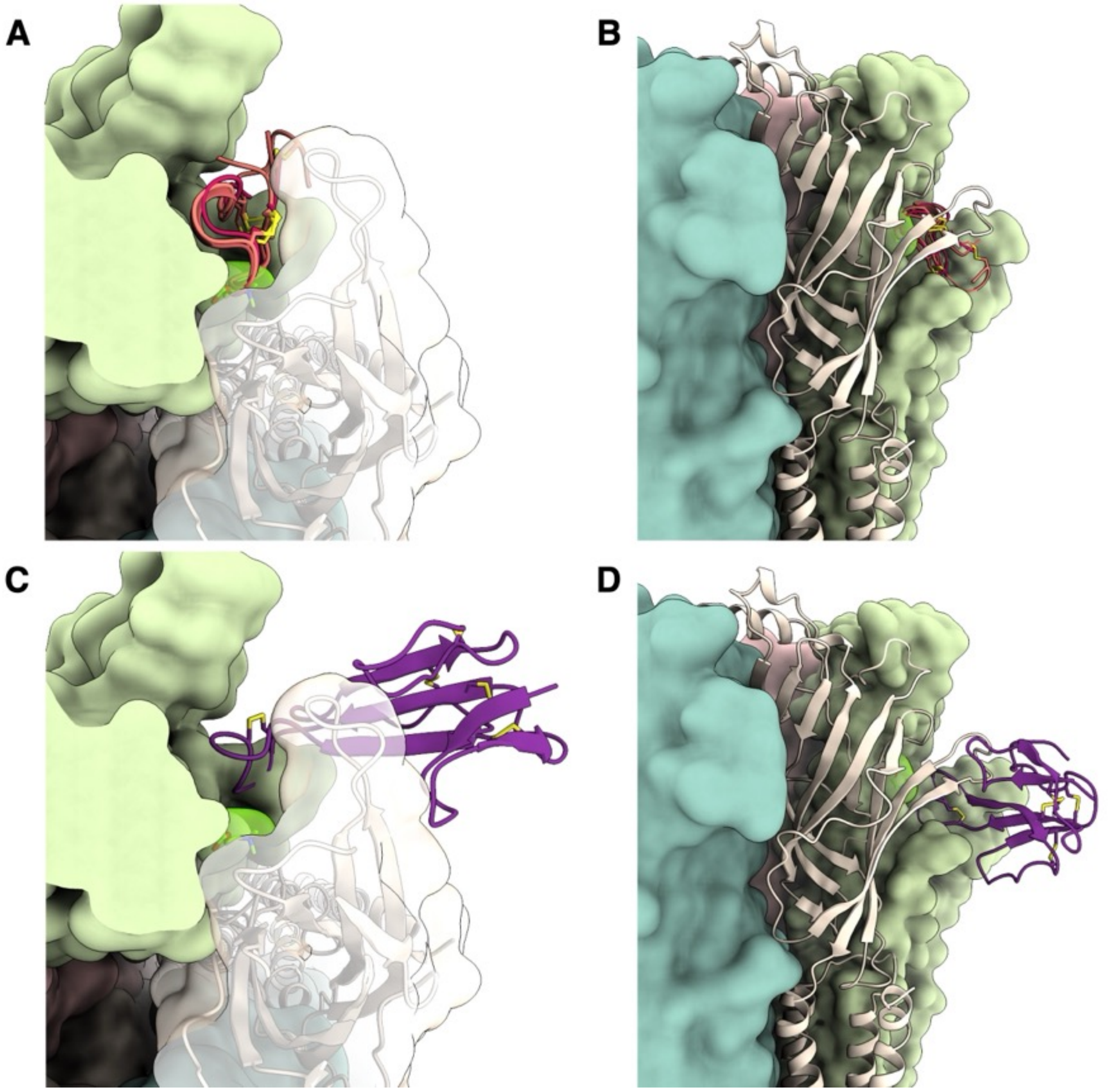

**Figure S8. Comparison of toxin binding at nAChR.**
(A and B) α-CTX used in this study seen from above and through the side between at the α-δ binding site. The ACh binding site is highlighted in lime green and overlaps with the proline-alanine residues of α-CTX. (C and D) Bungarotoxin is shown in purple in the same orientations as α-CTX.

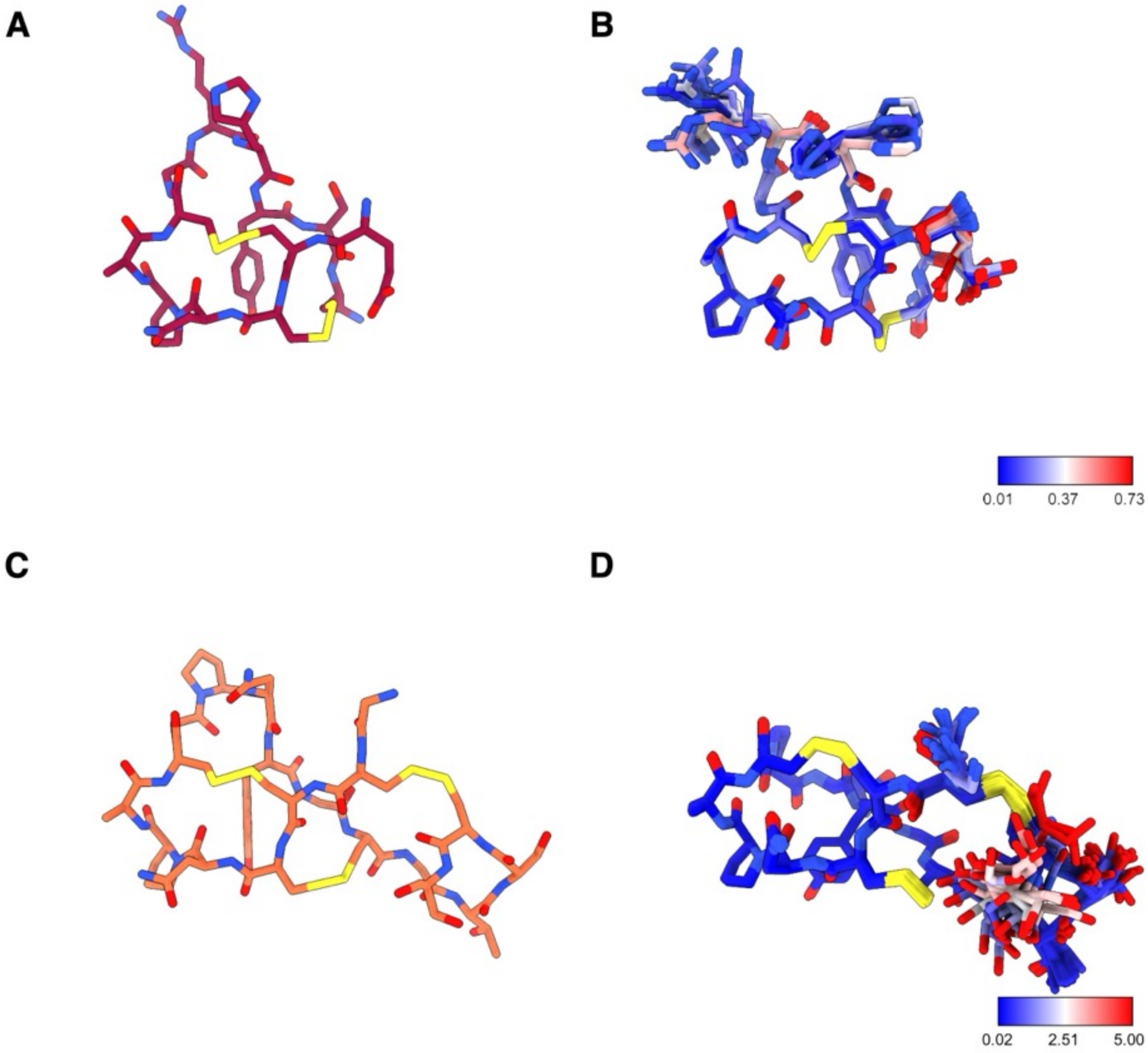

**Figure S9. Comparison of cryo-EM and NMR structures of α-CTX.**
(A,B) Cryo-EM structure of α-GI compared to the NMR structure (PDB 1xga). (C,D) Cryo-EM structure of α-SII compared to the NMR (PDB 6otb). Color legends around NMR display B-factors in Å$^2$.

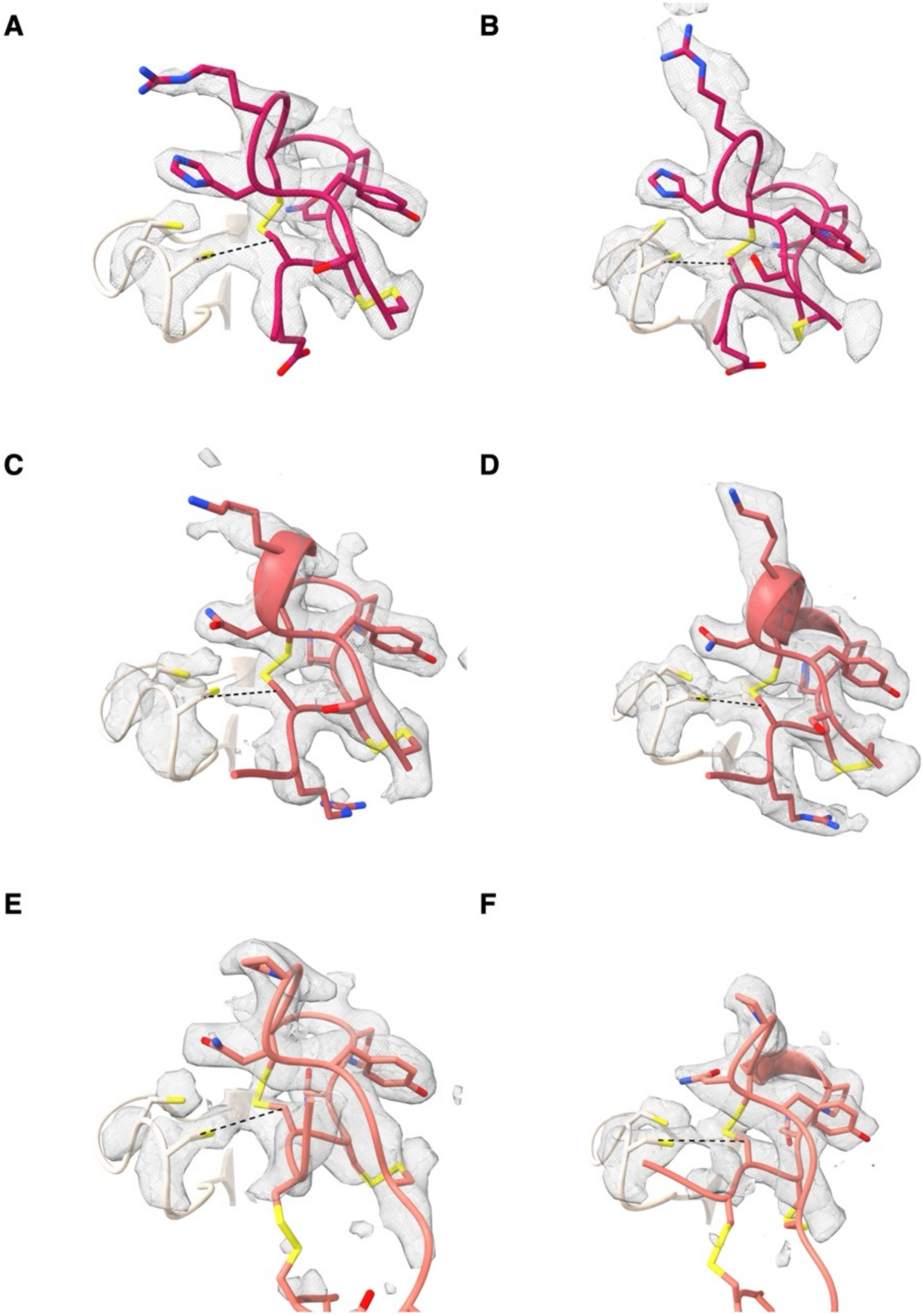

**Figure S10. Ambiguous density centered on the vicinal disulfide bond.**
Sharpened cryo-EM density maps for each toxin at their binding sites. (A,C,E) for the at α-δ and (B,D,F) at the α-ɣ. A,B α-GI, C,D α-MI. E,F α-SII. The dashed line connects αC192 and CII of the toxin.

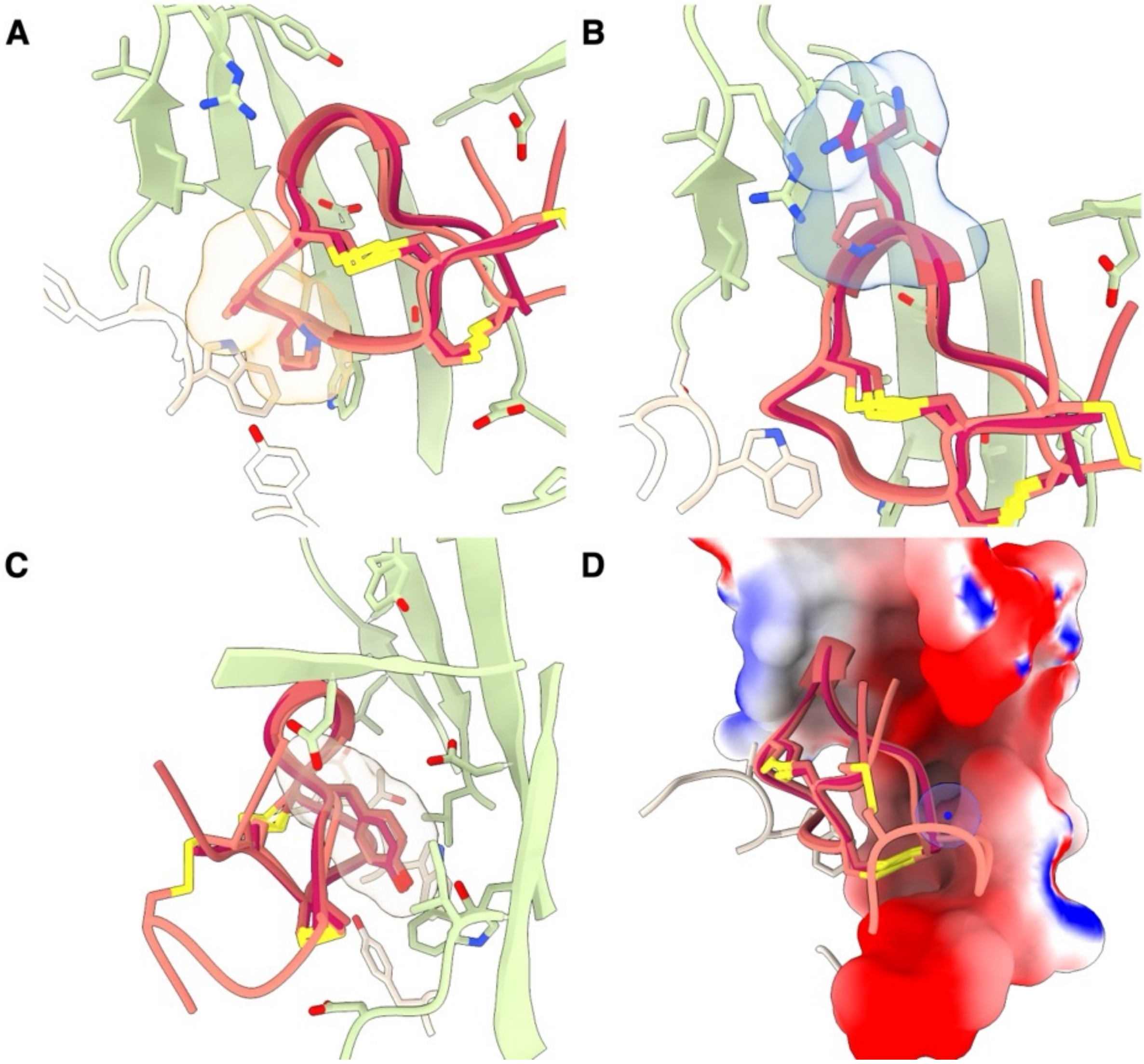

**Figure S11. Common features of α-CTX binding at the α-δ site.**
(A) The Pro-Ala lock at the cation binding site. (B) Cationic residue interacting with the complementary face. (C) Buried hydrophobic residue. (D) *C*'-terminal amide (highlighted blue sphere) with the acidic complementary face.

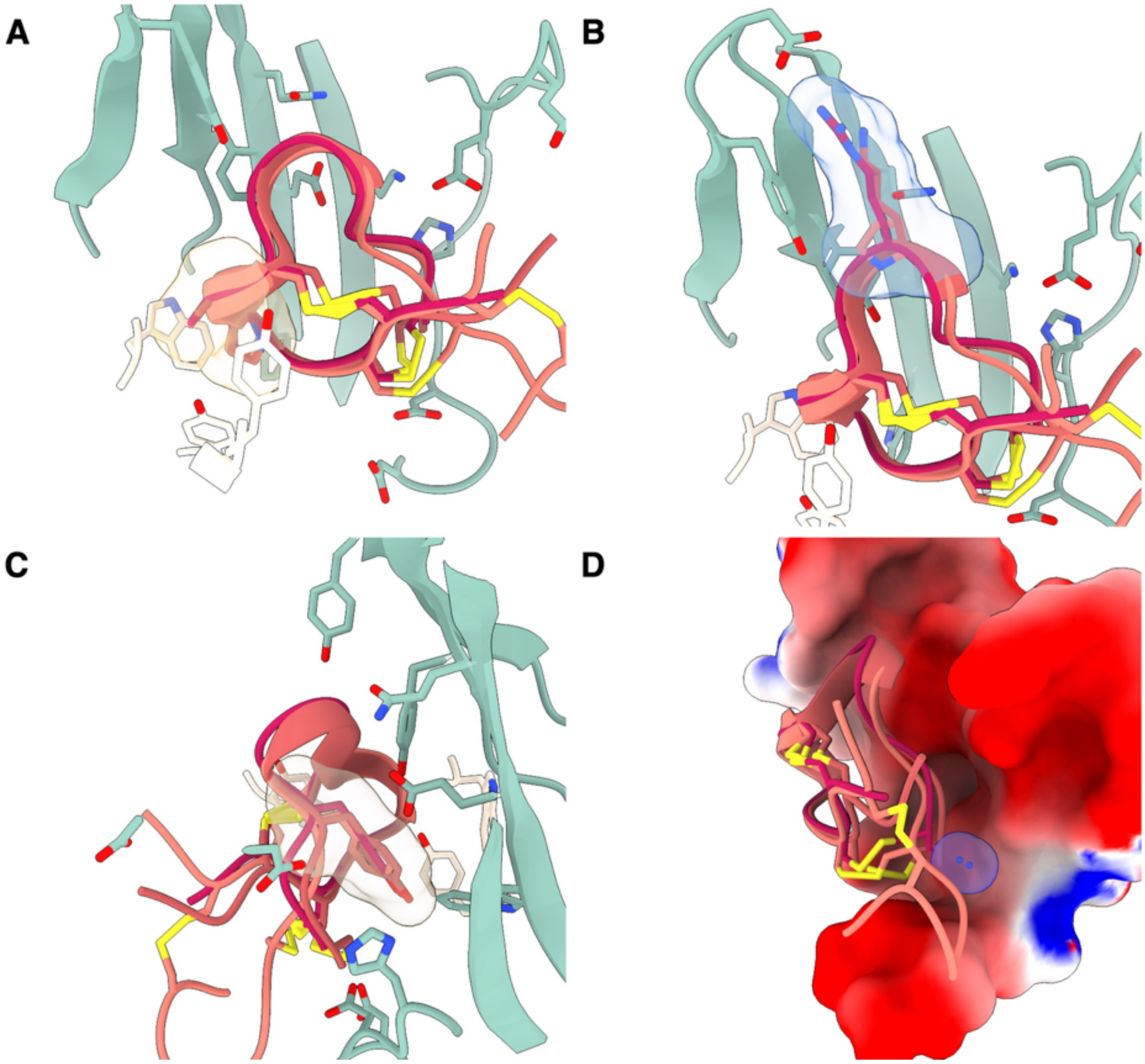

**Figure S12. Common eatures of α-CTX binding at the α-Ɣ site.**
(A) The Pro-Ala lock at the cation binding site. (B) Cationic residue interacting with the complementary face. (C) Buried hydrophobic residue. (D) *C*'-terminal amide (highlighted blue sphere) with the acidic complementary face.

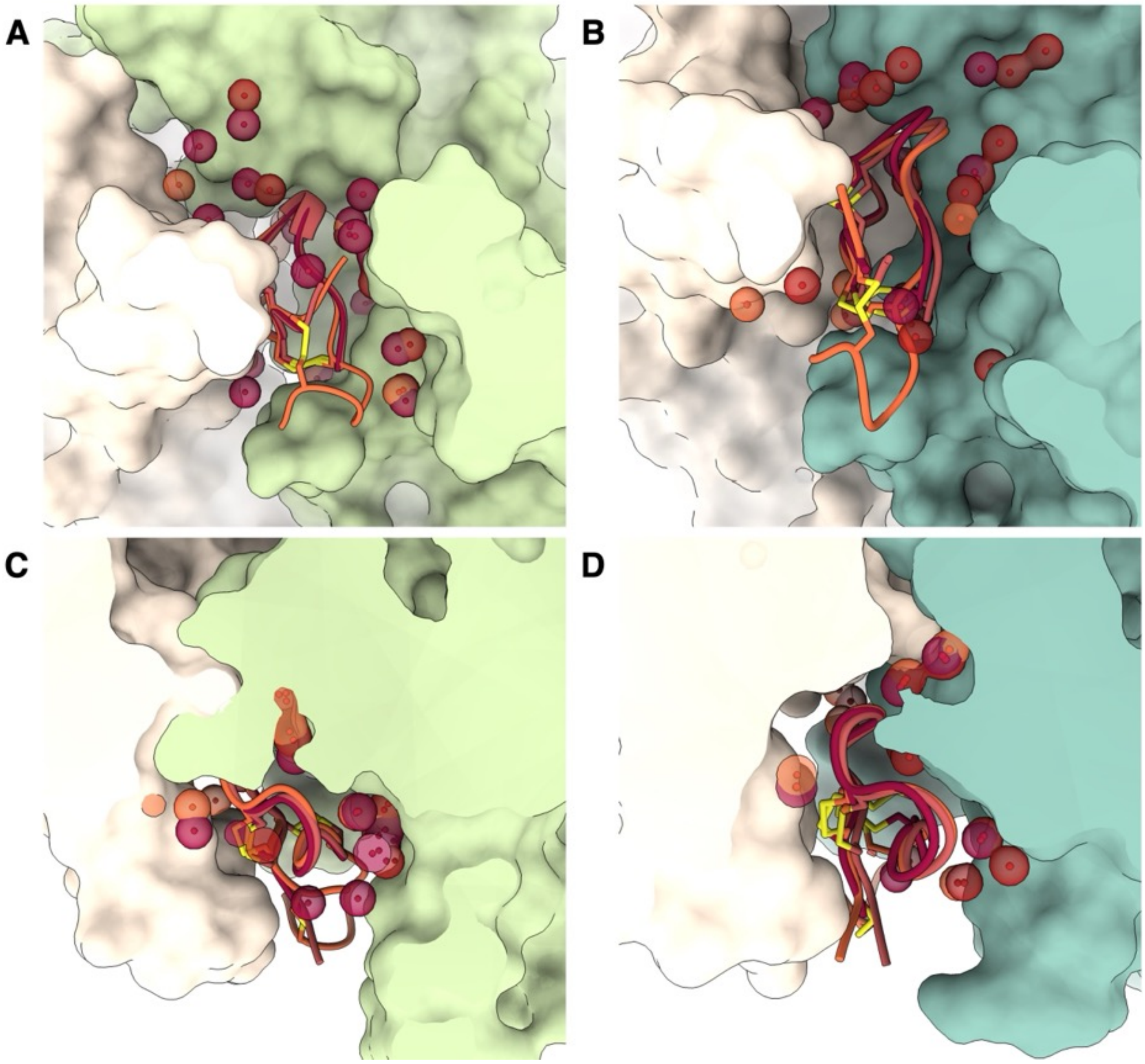

**Figure S13. Water present in the α-CTX binding sites**
Water determined by cryo-EM forming hydrogen bonds between the overlaid toxins and both the principal and complementary faces in red spheres. Alongside waters identified through computational water placement shown in orange.

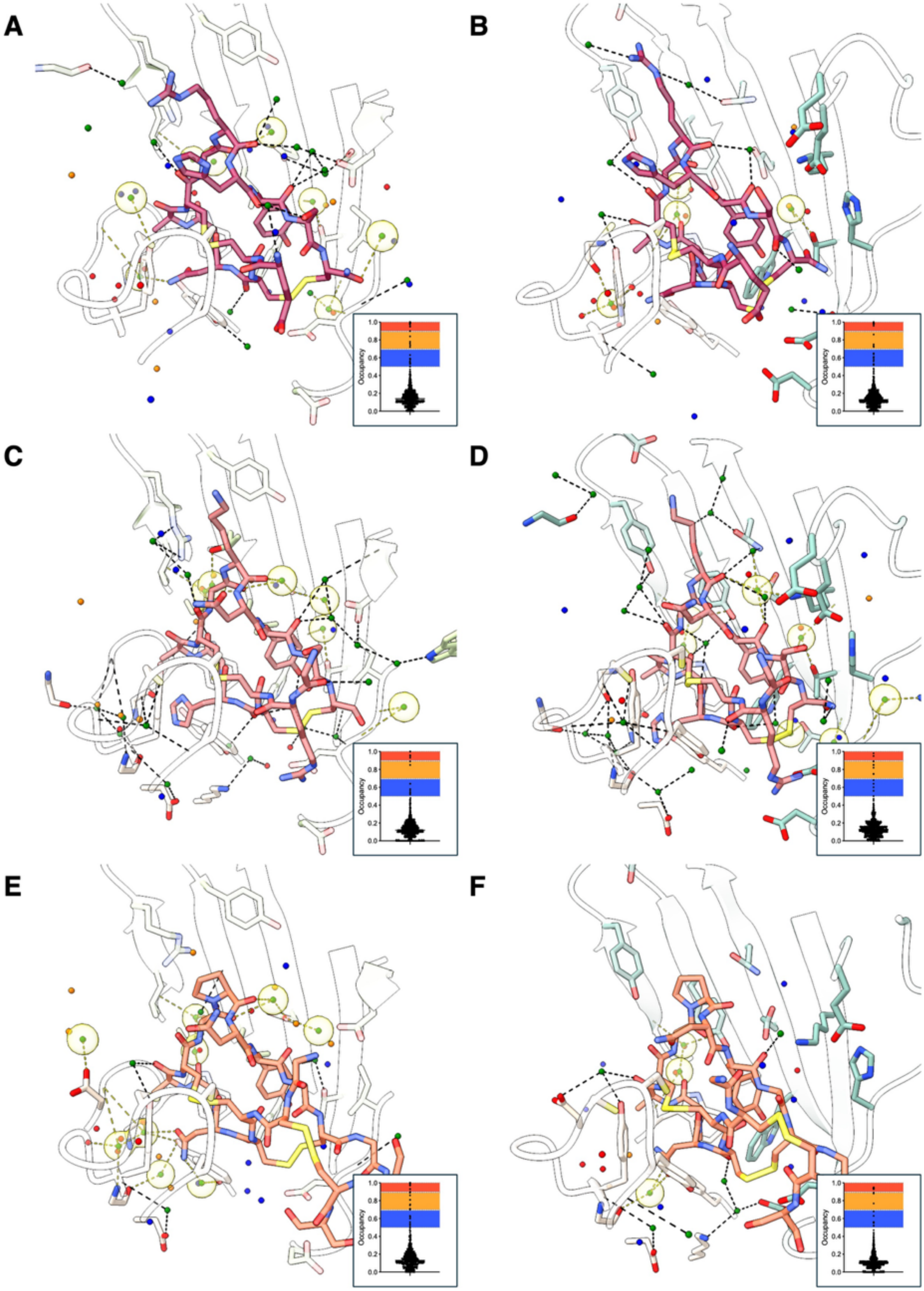

**Figure S14. Hydrogen bonding between waters identified in EM and confirmed by modelling.** Water identified at each binding site with cryo-EM water shown in green (A,C,E) for the at α-δ and (B,D,F) at the α-ɣ. A,B α-GI, C,D α-MI. E,F α-SII. Computational waters are shown as clusters in red, orange or blue according to confidence (violin plot distribution inset). Cryo-EM waters encased in yellow spheres are consistent with computational modelling.

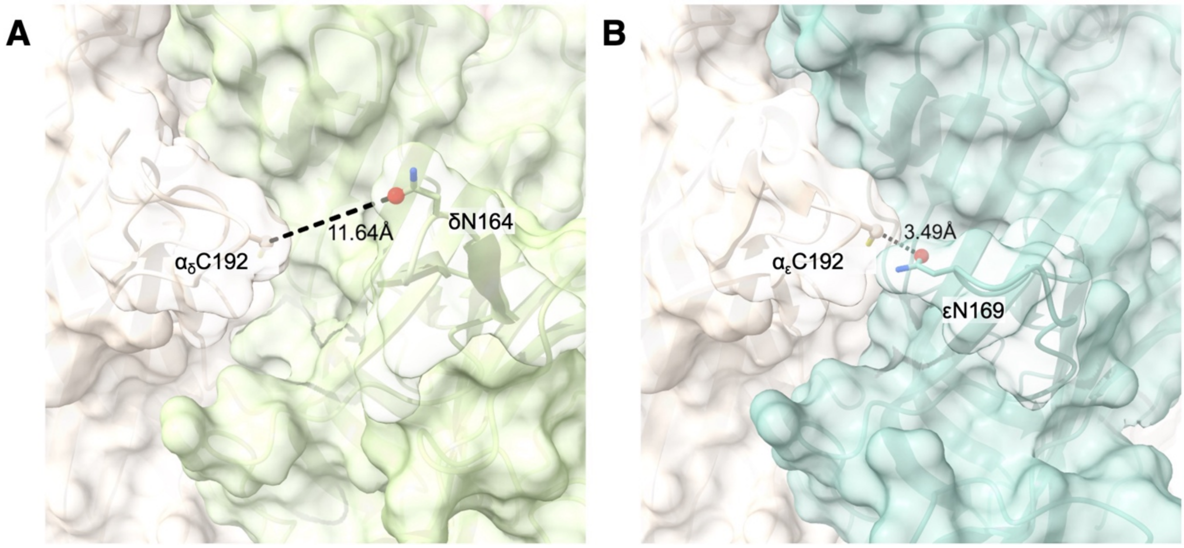

**Figure S15. Loop C and Loop F distances affect α-CTX docking at the human nAChR.** Computational docking proved intransigent at the α-γ site (B). The large discrepancy in the distance between αC192 and δN169 (11.6 Å) as shown in (A) compared to the equivalent distance between αC192 and γN164 of 3.5 Å.

### Additional peptide methods

#### *Table S5 Table of Peptides*

Name, sequence, % yield, % purity, *m/z* and retention time of peptides. Detailed characterisation data for peptides can be found in Figures **S16 – S25**.

| Peptide | Sequence | Yield (%) | % Purity | Calculated *m/z* | Observed *m/z* | $t_R$ (mins): gradient A | $t_R$ (mins): gradient B |
|---|---|---|---|---|---|---|---|
| **α-GI** | 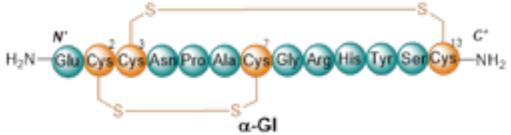  α-GI | 4 | 99 | 1437.61 | 1437.32 ± 0.08 | 10.3 | 18.7 |
| **α-MI** | 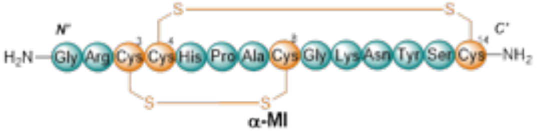  α-MI | 4 | 96 | 1494.57 | 1494.07 ± 0.47 | 14.3 | 20.1 |
| **α-SII** | 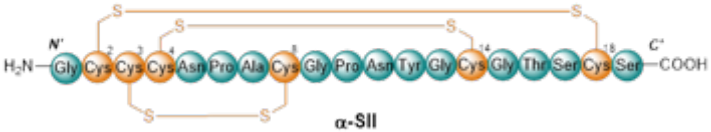  α-SII | 7 | 99 | 1790.97 | 1790.47 ± 0.50 | 11.0 | 18.7 |

### *Conotoxin Syntheses*
#### *Synthesis of Conotoxin α-GI*

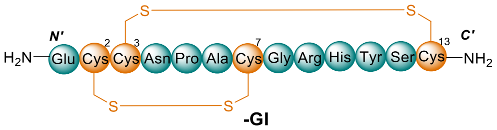

**-GI**

Linear α-GI was synthesised by automated Fmoc-SPPS as described in **General Protocol 1** (0.15 mmol). It was liberated from the resin and its protecting groups simultaneously removed under the conditions described in **General Protocol 3** to yield the crude linear α-GI as white amorphous powder (216 mg, 100% yield [based on initial resin loading]). Disulfide bond formation was facilitated in a solution of TFE/MQ $H_2O$ (1:1, *v/v*) as described by **General Protocol 4**. Cyclised α-GI was confirmed by analytical RP-HPLC and LCMS in moderate purity (68%). Solvent volume was initially reduced under vacuum prior to lyophilisation of the crude cyclised α-GI peptide.

Crude α-GI was solubilised in 0.1% TFA in MeCN:MQ $H_2O$ (2:8, *v/v*) at a concentration of ~40 mg/mL and purified by preparative RP-HPLC (3 x 1800 μL injection) employing a gradient of 5% – 65%B over 40 min (*ca.* 1.5%B/min) at a flow rate of 10 mL/min. Fractions were analysed by RP-HPLC and LCMS for compound identification and lyophilised to afford the compound, α-GI, as a white amorphous powder (9.3 mg, 99% purity, 4% overall yield).

**LCMS**: Mass calculated for [$C_{55}H_{80}N_{20}O_{18}S_4$ + H]: 1437.61; deconvoluted mass observed: 1437.32 ± 0.08. Charge states; 480.09 $[M+3H]^{3+}$, 719.64 $[M+2H]^{2+}$, 1438.41 $[M+H]^{+}$.

**RP-HPLC**: Phenomenex Aeris Peptide XB-C18 (100 Å, 5 µm, 150 mm x 4.6 mm), linear gradient A 5% – 95%B over 20 min (*ca.* 4.5%B/min) at 1 mL/min, $t_R$ = 10.3 min; linear gradient B 5% – 95%B over 50 min (*ca.* 1.8%B/min) at 1 mL/min, $t_R$ = 18.7 min.

#### *Synthesis of Conotoxin α-MI*

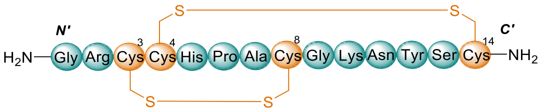

**-MI**

Linear α-MI was synthesised by automated Fmoc-SPPS as described in **General Protocol 1** (0.1 mmol). It was liberated from the resin and its protecting groups simultaneously removed under the conditions described in **General Protocol 3** to yield the crude linear α-MI as white amorphous powder (122 mg, 82% yield [based on initial resin loading]). Disulfide bond formation was facilitated in a solution of MeCN/MQ $H_2O$ (1:1, *v/v*) as described by **General Protocol 4**. Cyclised α-MI was confirmed by analytical RP-HPLC and LCMS in moderate purity (56%). Solvent volume was initially reduced under vacuum prior to lyophilisation of the crude cyclised α-MI peptide.

Crude α-MI was solubilised in 0.1% TFA in MeCN:MQ $H_2O$ (1:9, *v/v*) at a concentration of ~30 mg/mL and purified by preparative RP-HPLC (3 x 1400 μL injection) employing a gradient of 10% – 50%B over 50 min (*ca.* 0.8%B/min) at a flow rate of 10 mL/min. Fractions were analysed by RP-HPLC and LCMS for compound identification and lyophilised to afford the compound, α-MI, as a white amorphous powder (5.5 mg, 96% purity, 4% overall yield).

**LCMS**: Mass calculated for [$C_{55}H_{89}N_{22}O_{17}S_4$ + H]: 1494.57; deconvoluted mass observed: 1494.07 ± 0.47. Charge states; 499.08 $[M+3H]^{3+}$, 747.77 $[M+2H]^{2+}$, 1495.43 $[M+H]^{+}$.

**RP-HPLC**: Phenomenex Aeris Peptide XB-C18 (100 Å, 5 µm, 150 mm x 4.6 mm), linear gradient A 5% – 95%B over 60 min (*ca.* 1.5%B/min) at 1 mL/min, $t_R$ = 14.3 min; linear gradient B 5% – 50%B over 60 min (*ca.* 0.8%B/min) at 1 mL/min, $t_R$ = 20.1 min.

#### *Synthesis of Conotoxin α-SII*

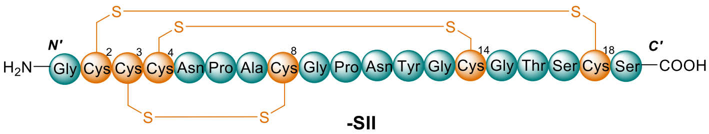

**-SII**

Linear α-SII was synthesised by manual Fmoc-SPPS as described in **General Protocol 2** (0.2 mmol). It was liberated from the resin and its protecting groups simultaneously removed under the conditions described in **General Protocol 3** to yield the crude linear α-SII as white amorphous powder (334 mg, 93% yield [based on initial resin loading]). Disulfide bond formation was facilitated in a solution of TFE/MQ $H_2O$ (1:1, *v/v*) as described by **General Protocol 4**. Cyclised α-SII was confirmed by analytical RP-HPLC and LCMS in moderate purity (50%). Solvent volume was initially reduced under vacuum prior to lyophilisation of the crude cyclised α-SII peptide.

Crude α-SII was solubilised in 0.1% TFA in MeCN:MQ $H_2O$ (2:8, *v/v*) at a concentration of ~40 mg/mL and purified by preparative RP-HPLC (5 x 1800 μL injection) employing a gradient of 5% – 65%B over 40 min (*ca.* 1.5%B/min) at a flow rate of 10 mL/min. Fractions were analysed by RP-HPLC and LCMS for compound identification and lyophilised to afford the compound, α-SII, as a white amorphous powder (24.1 mg, 99% purity, 7% overall yield).
**LCMS**: Mass calculated for [$C_{66}H_{95}N_{21}O_{26}S_6$ + H]: 1790.97; deconvoluted mass observed: 1790.47 ± 0.50. Charge states; 896.41$[M+2H]^{2+}$, 1791.12 $[M+H]^{+}$.
**RP-HPLC**: Phenomenex Aeris Peptide XB-C18 (100 Å, 5 μm, 150 mm x 4.6 mm), linear gradient A 5% – 95%B over 20 min (*ca.* 4.5%B/min) at 1 mL/min, $t_R$ = 11.0 min; linear gradient B 5% – 95%B over 50 min (*ca.* 1.8%B/min) at 1 mL/min, $t_R$ = 18.7 min.

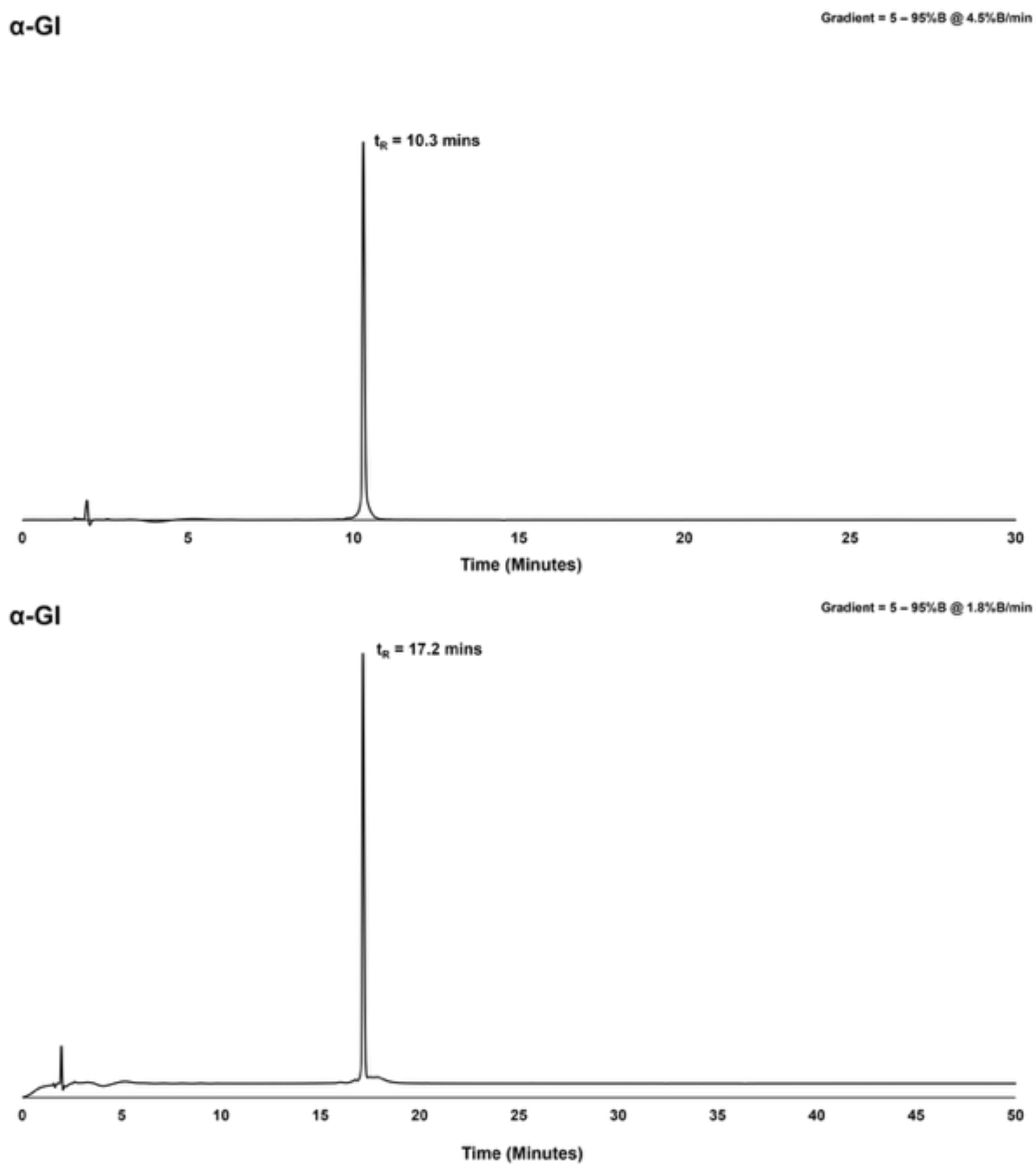

***Figure S16*** Analytical RP-HPLC chromatogram (214 nm) of purified peptide, **α-GI** (*ca.* 99% as analysed by peak area. Phenomenex Aeris Peptide XB-C18 (100 Å, 5 µm, 150 mm x 4.6 mm), **upper:** linear gradient A 5% – 95%B over 20 min (*ca.* 4.5%B/min) at 1 mL/min, $t_R$ = 10.3 min; **lower:** linear gradient B 5% – 95%B over 50 min (*ca.* 1.8%B/min) at 1 mL/min, $t_R$ = 18.7 min.

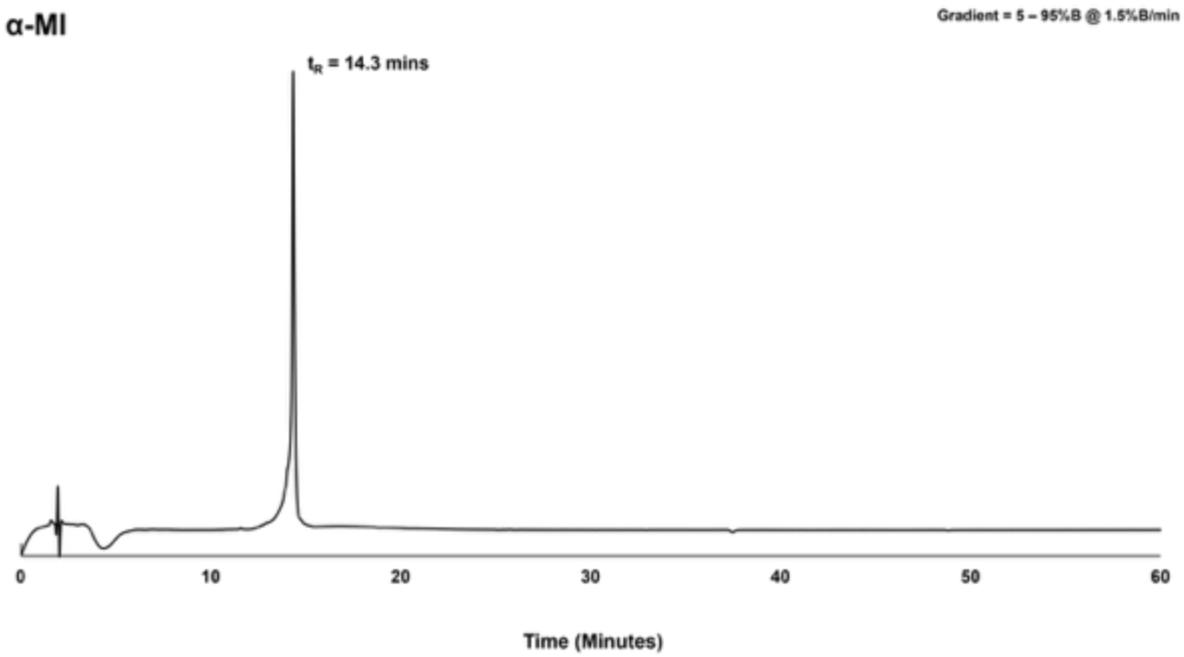

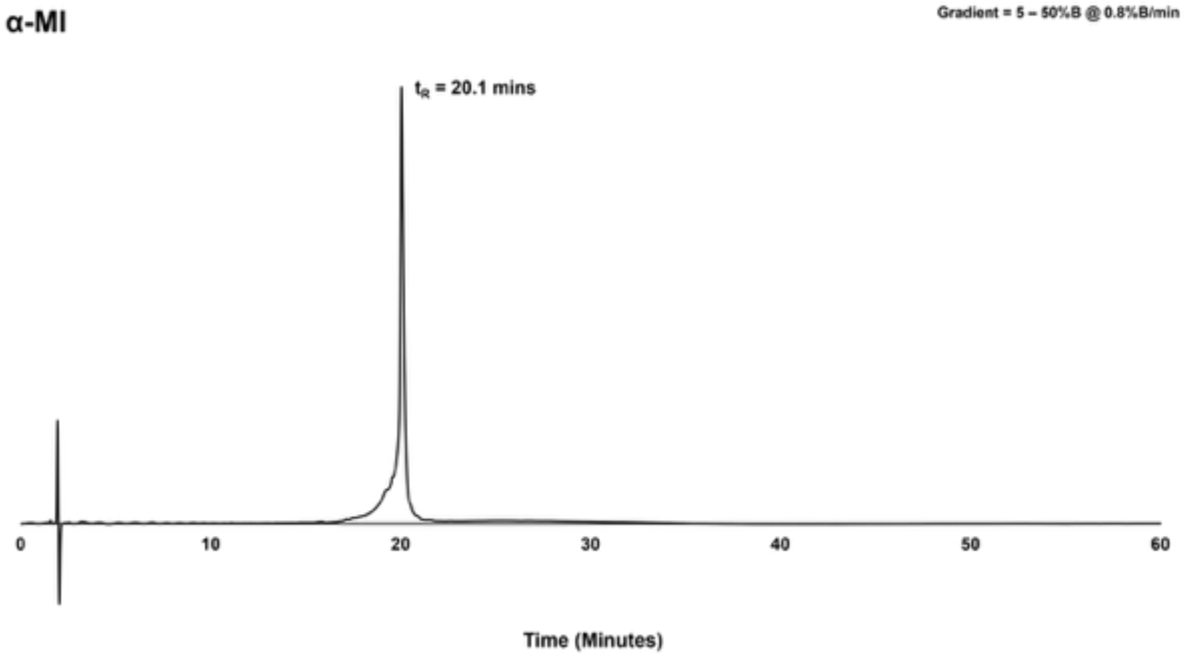

***Figure S17*** Analytical RP-HPLC chromatogram (214 nm) of purified peptide, **α-MI** (*ca.* 96% as analysed by peak area. Phenomenex Aeris Peptide XB-C18 (100 Å, 5 µm, 150 mm x 4.6 mm), **upper:** linear gradient A 5% – 95%B over 60 min (*ca.* 1.5%B/min) at 1 mL/min, $t_R$ = 14.3 min; **lower:** linear gradient B 5% – 50%B over 60 min (*ca.* 0.8%B/min) at 1 mL/min, $t_R$ = 20.1 min.

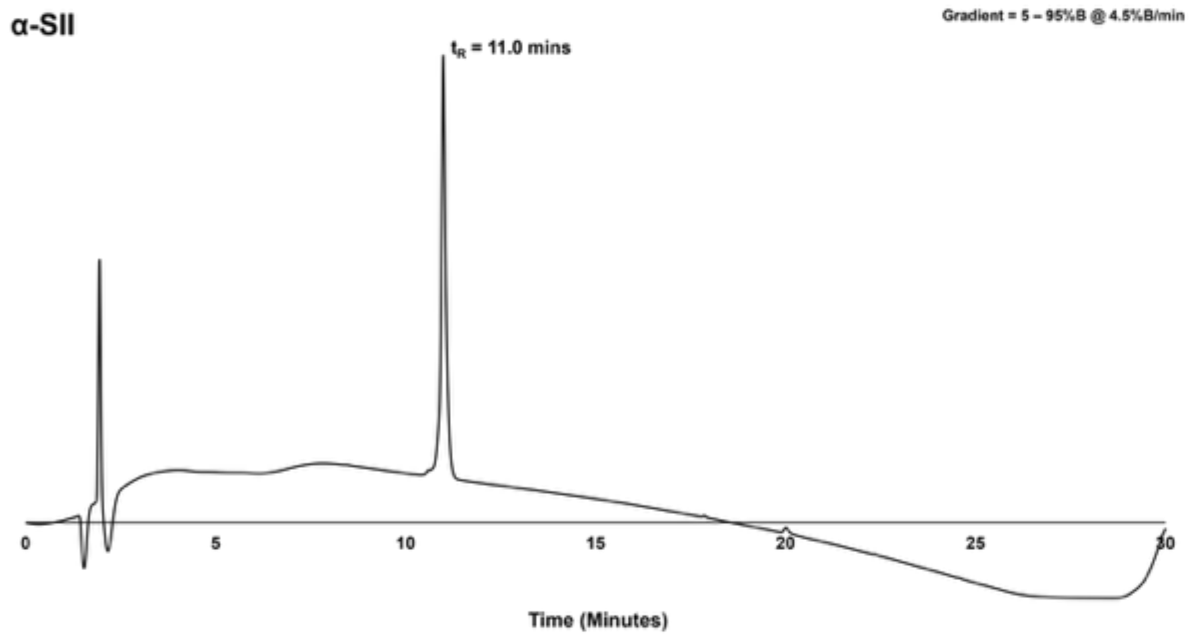

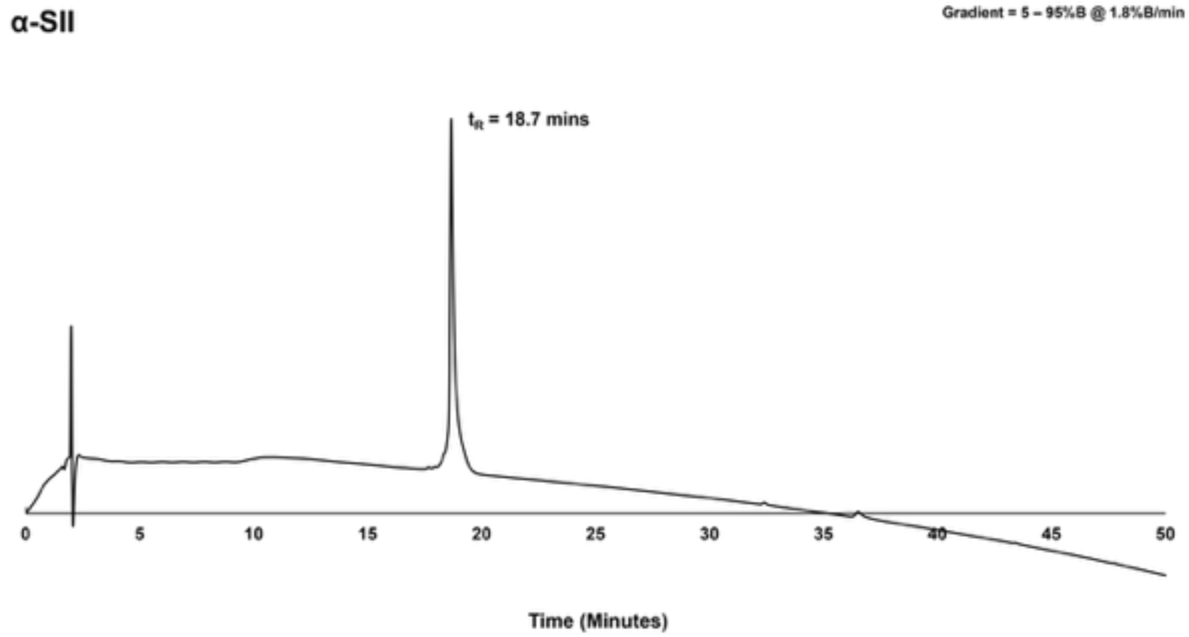

***Figure S18*** Analytical RP-HPLC chromatogram (214 nm) of purified peptide, **α-SII** (*ca.* 99% as analysed by peak area. Phenomenex Aeris Peptide XB-C18 (100 Å, 5 µm, 150 mm x 4.6 mm), **upper:** linear gradient A 5% – 95%B over 20 min (*ca.* 4.5%B/min) at 1 mL/min, $t_R$ = 11.0 min; **lower:** linear gradient B 5% – 95%B over 50 min (*ca.* 1.8%B/min) at 1 mL/min, $t_R$ = 18.7 min.

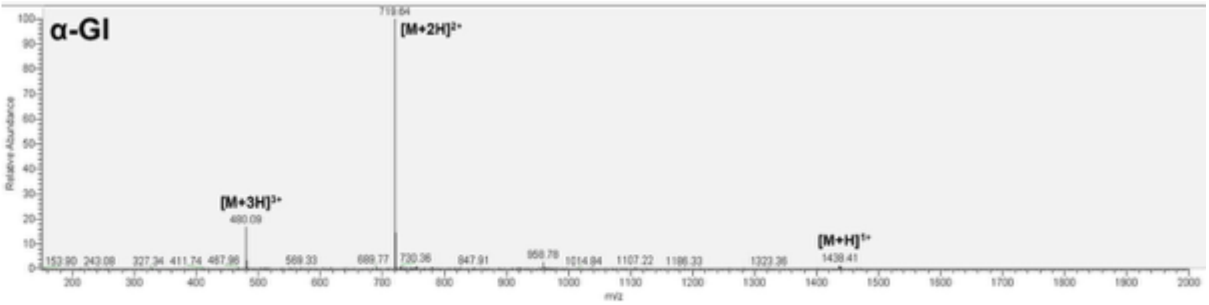

***Figure S19*** LCMS of purified **α-GI**, mass calculated for [$C_{55}H_{80}N_{20}O_{18}S_4$ + H]: 1437.61; deconvoluted mass observed: 1437.32 ± 0.08. Charge states; 480.09 $[M+3H]^{3+}$, 719.64 $[M+2H]^{2+}$, 1438.41 $[M+H]^{+}$.

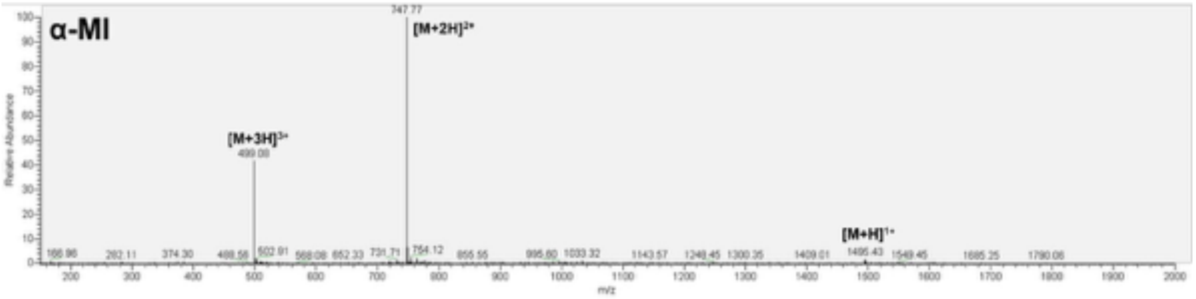

***Figure S20*** LCMS of purified **α-MI**, mass calculated for [$C_{55}H_{89}N_{22}O_{17}S_4$ + H]: 1494.57; deconvoluted mass observed: 1494.07 ± 0.47. Charge states; 499.08 $[M+3H]^{3+}$, 747.77 $[M+2H]^{2+}$, 1495.43 $[M+H]^{+}$.

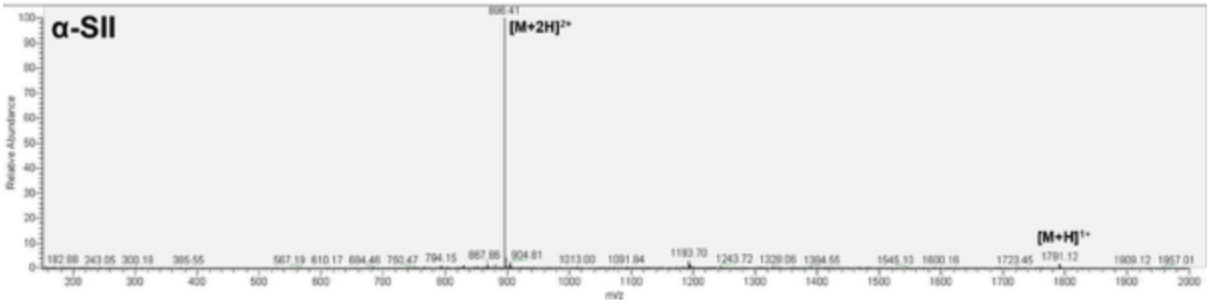

***Figure S21*** LCMS of purified **α-SII**, mass calculated for [$C_{66}H_{95}N_{21}O_{26}S_6$ + H]: 1790.97; deconvoluted mass observed: 1790.47 ± 0.50. Charge states; 896.41 $[M+2H]^{2+}$, 1791.12 $[M+H]^{+}$.

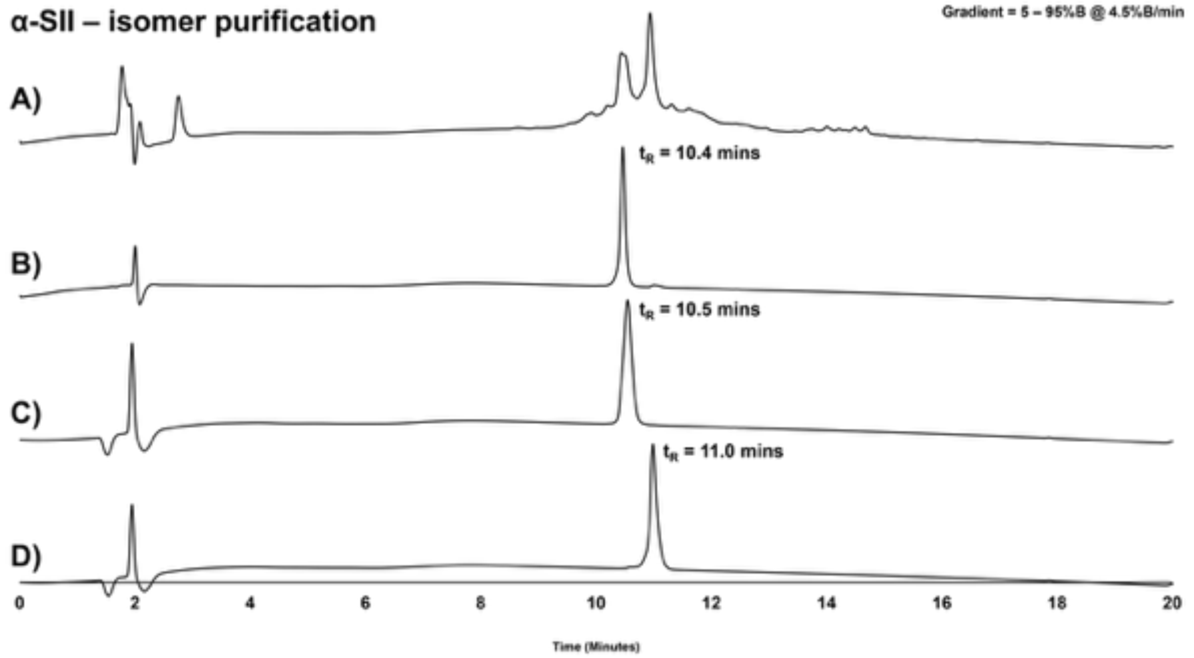

*Figure S22* Analytical RP-HPLC chromatograms (214 nm) for the separation of isomers of α-SII. A) Crude peptide post oxidation. B) Isomer 1. C) Isomer 2. D) Isomer 3 (desired isomer as confirmed by activity and Cryo-EM).

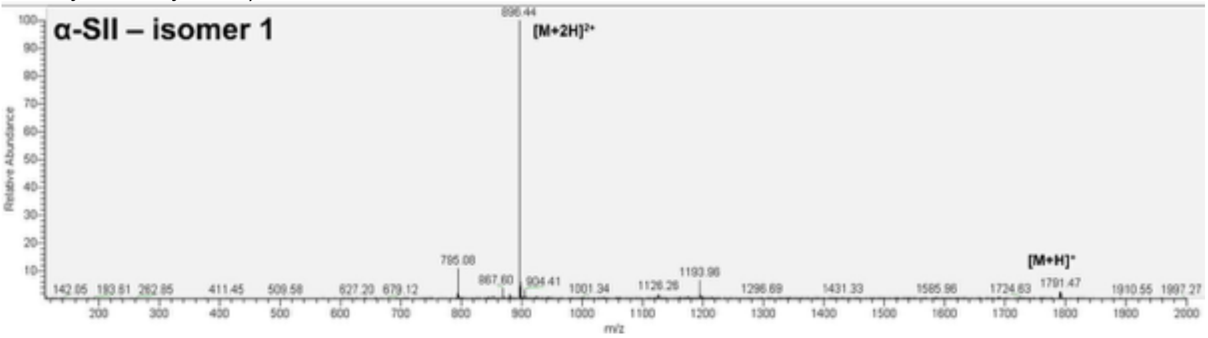

*Figure 23* LCMS of purified isomer 1 of **α-SII** corresponding to peak at $t_R$ = 10.4 mins in **Figure S22**, mass calculated for $[C_{66}H_{95}N_{21}O_{26}S_6 + H]$: 1790.97; deconvoluted mass observed: 1790.68 ± 0.29. Charge states; 896.44 $[M+2H]^{2+}$, 1791.47 $[M+H]^+$.

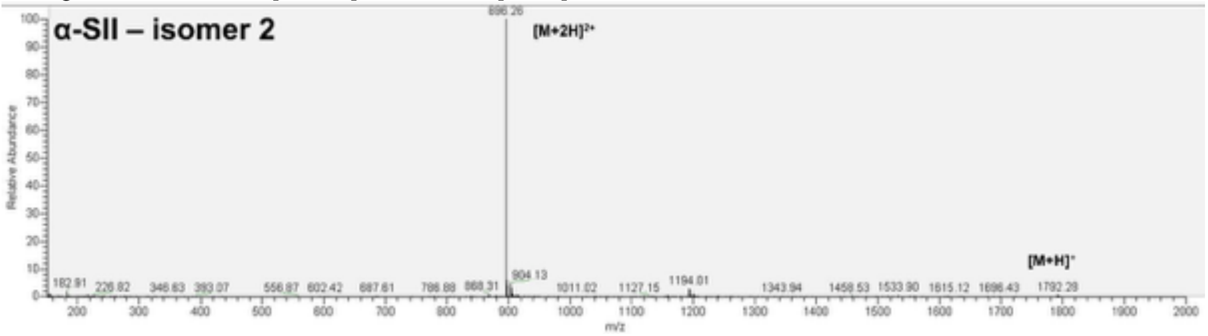

*Figure S24* LCMS of purified isomer 2 of **α-SII** corresponding to peak at $t_R$ = 10.5 mins in **Figure S22**, mass calculated for $[C_{66}H_{95}N_{21}O_{26}S_6 + H]$: 1790.97; deconvoluted mass observed: 1790.40 ± 0.17. Charge states; 896.26 $[M+2H]^{2+}$, 1791.28 $[M+H]^+$.

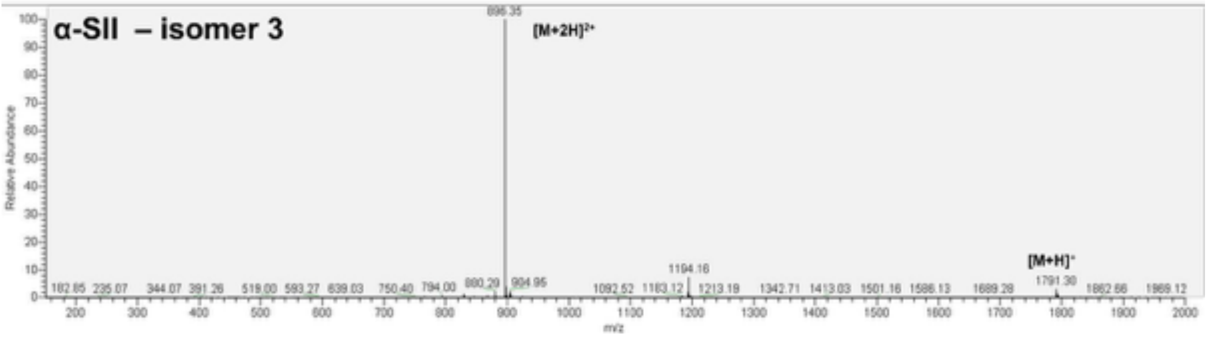

*Figure S25* LCMS of purified isomer 3 of **α-SII** corresponding to peak at $t_R$ = 11.0 mins in **Figure S22**, mass calculated for [$C_{66}H_{95}N_{21}O_{26}S_6$ + H]: 1790.97; deconvoluted mass observed: 1790.50 ± 0.28. Charge states; 896.35 $[M+2H]^{2+}$, 1791.30 $[M+H]^{+}$.
